# Chromatin Perturbation Promotes Susceptibility to Hypomethylating Agents

**DOI:** 10.1101/2025.07.28.666975

**Authors:** Constanze Schneider, Gabriela Alexe, Lucy A. Merickel, Ashleigh Meyer, Michelle L. Swift, Rodolfo B. Serafim, Allen T. Basanthakumar, Audrey Taillon, Silvi Salhotra, Fabio Boniolo, Sander Lambo, Yara Rodriguez, Björn Häupl, Rani E. George, David E. Root, Thomas Oellerich, Volker Hovestadt, Dipanjan Chowdhury, Kimberly Stegmaier

## Abstract

Cancer-directed drugs are often clinically deployed without definitive understanding of their molecular mechanisms of action (MOA). Hypomethylating agents (HMAs), which result in the degradation of the DNA methyltransferase 1 (DNMT1), have been deployed for decades in the treatment of haematological malignancies^1,2^. The precise mechanism of action of these drugs, however, has been debated, rendering the design of rational combination therapies challenging. Here, we identified the deubiquitinating enzyme USP48 as a crucial regulator of posttranslational histone modification in the context of DNA demethylation. USP48 loss selectively enhances response to DNMT1 inhibition, leading to a rapid induction of cell death. We demonstrate that USP48 is localized at sites of DNA damage and deubiquitinates H2A variants and proteins important for DNA damage repair. Functionally, loss of USP48 triggers an increase in chromatin accessibility upon HMA treatment, rendering AML cells more susceptible to DNA damage. Our results support USP48 as a posttranslational histone modifier for chromatin stability and DNA damage in response to HMA-related DNA demethylation. These findings propose USP48 as a new target for combination therapy with HMAs for acute myeloid leukaemia (AML).

---

Most traditional cytotoxic chemotherapies do not inhibit singular targets but rather broadly affect multiple pathways, making understanding of their clinically relevant mechanisms of action (MOA) challenging. Even targeted therapies, such as selective kinase inhibitors, show unintended off-target effects^3^. Gaining a better understanding of how this polypharmacology contributes to anti-cancer effects observed in patients could reveal potential vulnerabilities and novel combination strategies to enhance cancer therapies.

Hypomethylating agents (HMAs), such as azacitidine (AZA) and decitabine (DAC), are used in patients with myelodysplastic syndrome (MDS) and acute myeloid leukaemia (AML) who are not eligible for intensive chemotherapy^1,2^. The active metabolites of both drugs, analogues of the nucleoside cytidine, are incorporated into DNA and irreversibly bind DNA methyltransferase 1 (DNMT1)^4^, inducing its proteasomal degradation. The downstream consequences of HMA treatment are multi-faceted and include 1) DNA damage, caused by DNA-DNMT1 protein adducts^5^, 2) altered chromatin integrity, due to loss of epigenetic silencing of regions in the genome^6,7^, 3) demethylation of gene promoters, silenced during cancer development, and 4) reactivation of endogenous retroviral elements, triggering the activation of an inflammatory response in the cell^8^. It is not clear which of these mechanisms are contributing to clinical responses, and thus rational combinations based on MOA have been limited. New chemistry specifically inhibiting the enzymatic activity of DNMT1 coupled with unbiased screening approaches enable identification of rational combinations exploiting the hypomethylating capacity of these drugs. Using the potent first-in-class DNMT1-selective inhibitor GSK-3685032^9^ allows for the mechanistic dissection of the effects of DNA demethylation on AML biology, bypassing the multi-faceted MOA of previous deployed HMAs.

Herein, we aimed to identify active combinations with HMAs which can be therapeutically exploited, utilizing unbiased CRISPR screens, proteogenomic approaches, and the novel non-nucleoside analogue DNMT1 inhibitor GSK-3685032^9^. Our integrated analysis of clinically used HMAs with GSK-3685032 identified a selective dependency of AML cells on the deubiquitinating enzyme USP48 upon DNMT1 inhibition, presenting a tractable strategy for the development of potent combination therapies and the prospect to enhance patient outcome.

## HMA-specific sensitizer screen

Using an unbiased genome-wide screening approach, we aimed to identify novel targets sensitizing AML cells to HMAs. Therefore, MV4-11 cells were transduced with Cas9 and the genome-scale Avana sgRNA library and cultured for 16 days under azacytidine (AZA), decitabine (DAC) or GSK-3685032 (GSK) treatment versus a DMSO control (Fig. 1a). Due to the potential impact of HMAs on both DNA damage through their incorporation into DNA and on DNA methylation, drug concentrations were picked to achieve an IC30 or IC10 for the nucleoside analogue drugs (AZA and DAC). The DNMT1-selective inhibitor (GSK-3685032) was included because it induces hypomethylation without direct DNA damage through adduct formation. Hits enriched in AZA and DAC showed a strong overlap (Fig. 1b), consistent with their overlapping mechanisms of action. Genes previously associated with drug resistance, such as the nucleoside transporter *SLC29A1* and the kinases *UCK2* (AZA) and *DCK* (DAC), important for mono-phosphorylation of the inactive metabolites, scored strongly for resistance in our screens (Fig. 1b and c)^10^. Furthermore, our results validated previous findings that the knockout of the dNTP hydrolase *SAMHD1* sensitizes to DAC, but not AZA ^11^. *TOPORS*, encoding an E3 ubiquitin ligase related to resolving DNMT1-DNA crosslinks and DNA damage repair (DDR), which scores as a sensitizer solely in the HMA higher concentration, was also previously linked to HMA response in AML (Fig. 1c)^12,13^.

**Fig. 1:**
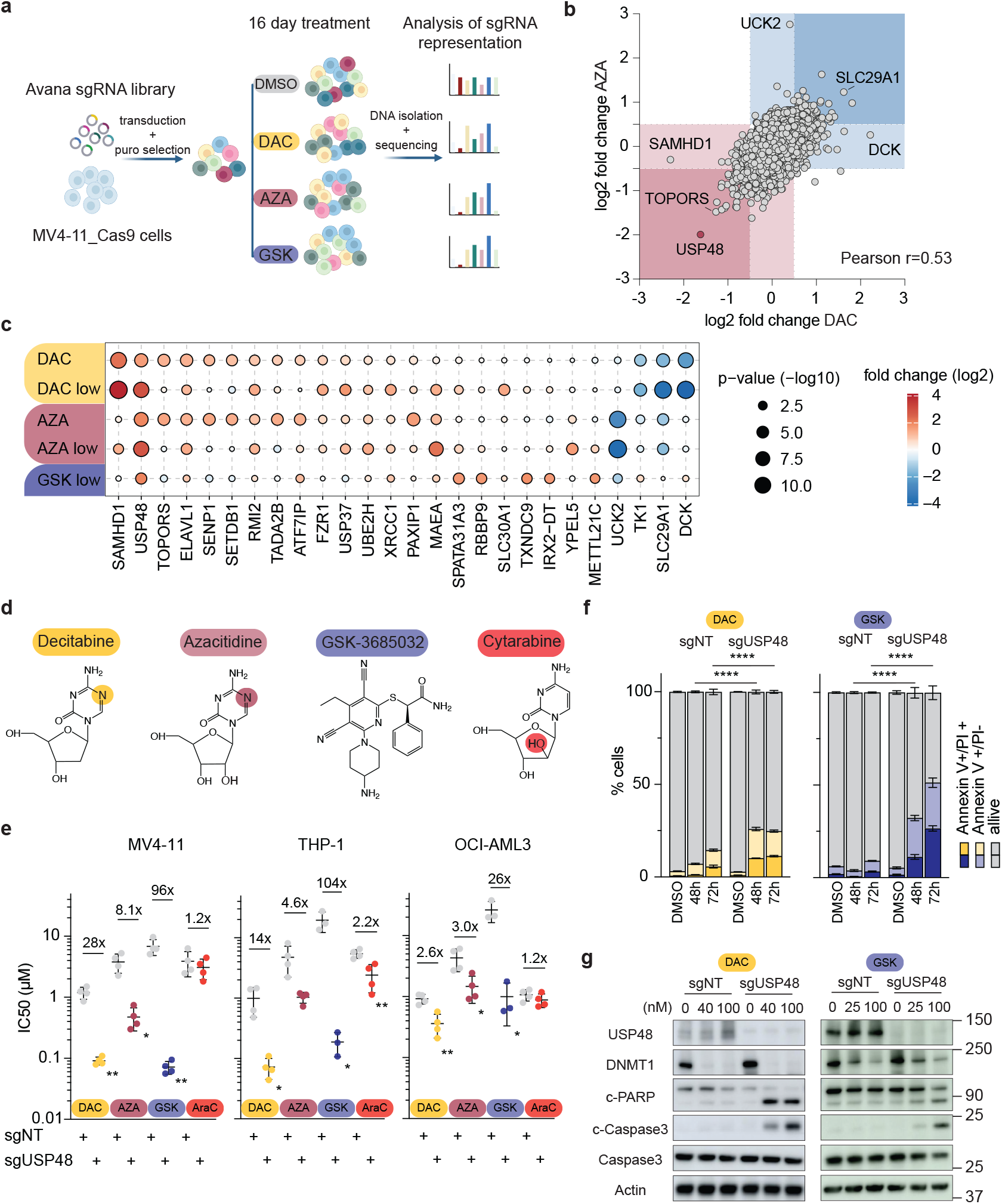
HMA-sensitizer screen identified USP48 as target in AML. **a)** Schematic overview of the CRISPR/Cas9 screening approach with azacitidine (AZA), decitabine (DAC) or GSK-3685032 (GSK) treatment and DMSO as a control (created with BioRender). **b)** Scatter dot plot showing the log2 fold change (LFC) of sgRNA abundance in decitabine (DAC) treatment vs. DMSO and azacitidine (AZA) treatment vs. DMSO. The scores of all sgRNAs targeting the same gene were averaged and shown as one data point, with resistance hits highlighted in blue and sensitizer hits in red. Significance cutoff: hypergeometric test abs (log2 fold change) ≥ 0.5, average p-value ≤ 0.1. **c)** Bubble plot of the top 22 sensitizer hits and 4 resistant hits enriched in the treatment arms. Bubble size represents the p-value, colour indicates the average log2 fold change per target. **d)** Compound structures of the nucleoside analogue HMAs decitabine and azacytidine, the DNMT1 inhibitor GSK-3685032 and the nucleoside analogue drug cytarabine. The modifications in comparison to deoxycytidine are colour coded. **e)** Dot plot representing the IC50 values for MV4-11, THP-1 and OCI-AML3 carrying sgNT (grey) or sgUSP48 (coloured) for DAC (yellow), AZA (burgundy), GSK-3685032 (blue) and AraC (red). Viability was assessed after 96h (n=4). 2-way ANOVA multiple comparisons test was used to compare treatment conditions in sgUSP48 vs. sgNT per cell line * p < 0.05, ** p < 0.01. **f)** Annexin V/PI staining in MV4-11 sgNT and sgUSP48 cells treated with 100 nM DAC (yellow) or 20 nM GSK-3680532 (blue) for 48 and 72h. Percent of early apoptotic (Annexin V+/PI-) or late apoptotic (Annexin V+/PI+) cells are shown per condition (n=3). One-way ANOVA multiple comparisons test with Tukey’s correction was used to compare sgUSP48 treated vs. sgNT treated conditions per time point **** p < 0.0001. **g)** Western blot validation of *USP48* KO and levels of DNMT1, cleaved PARP and cleaved Caspase 3 in MV4-11 sgNT and sgUSP48 cells treated with multiple concentrations of decitabine or GSK-3680532 for 48h. Actin serves as loading control.

Functional clustering of the core 300 overlapping sensitizers showed an enrichment of chromatin modifiers, immune response genes, DNA damage repair (DDR) related targets, and Ubiquitin/SUMO-ylating enzymes for all DNMT1 inhibitors (Extended Data Fig. 1a and b, Extended Data Table 1). Loss of genes related to OXPHOS and cell cycle led to enhanced resistance in all conditions, presumably by affecting proliferation and therefore the susceptibility to DNMT1 inhibitory effects (Extended Data Fig. 1c)^14^. While GSK-3685032 shared many of the same gene set enrichments as the other HMAs, a subset of targets related to MAPK and Rho-GTPases signalling was found specifically in the GSK-3685032 treatment arm (Extended Data Fig. 1d).

USP48, a deubiquitinating enzyme, scored strongest in all HMA conditions and was not a dependency in the DMSO control (Extended Data Fig. 1e). Analysis of the Cancer Dependency Map (DepMap) data confirmed that USP48 does not score as a strong dependency (gene effect score < -0.5) in any subtype or lineage (Extended Data Fig. 1f). Cancer Cell Line Encyclopedia (CCLE) expression data, as well as expression data from multiple patient cohorts, showed a universally high expression of *USP48* with increased expression in AML compared to other cancer types (Extended Data Fig. 1g). USP48 has been described to stabilize proteins like gasdermin E, aurora B or NF-κB/p65, and influence multiple pathways in different disease settings^15–18^. Literature on USP48, however, is sparse and does not explore its function in AML or HMA treatment response.

### USP48 loss induces cell death upon HMA treatment

To explore the role of USP48 in AML, we deployed a doxycycline inducible CRISPR/Cas9 system targeting *USP48* and a non-targeting guide as a control (Extended Data Fig. 2a). Low-throughput validation in multiple AML cell lines was performed, testing the two structurally related nucleoside analogue-based HMAs (DAC and AZA), the DNMT1 inhibitor (GSK-3685032), and the nucleoside analogue drug cytarabine (AraC) (Fig. 1d). MV4-11 cells carrying sgRNAs against *USP48* showed a significant increase in HMA toxicity (DAC: 28-fold, AZA: 8-fold and GSK: 96-fold) in comparison to cells carrying the non-targeting control guide (Fig. 1e, Extended Data Fig. 2b). In contrast, no differential response (1.2-fold) was detected in the cytarabine treated condition (Fig. 1e, Extended Data Fig. 2b). Similar results were observed in THP-1 and OCI-AML3 cells, confirming the significant shift in HMA sensitivity, while other nucleoside analogue drugs such as AraC, DNA damage-inducing drugs including doxorubicin, the topoisomerase I inhibitor SN-38, and the PARP inhibitor talazoparib, largely showed no increased activity upon *USP48* knockout (KO) (Fig. 1e, Extended Data Fig. 2c and d). These data suggest that the effect of *USP48* loss is specifically linked to DNA hypomethylation and is independent of the incorporation of nucleoside analogues into the DNA or the induction of DNA damage through generation of DNA adducts.

To assess which mechanism is promoting alteration in cell viability in *USP48* KO cells in combination with HMA treatment, we evaluated the induction of differentiation, cell cycle arrest or cell death. Only modest changes were observed in cell morphology using May-Grünwald-Giemsa staining or expression of CD11b, a surface marker for monocytic differentiation, upon DAC treatment +/-*USP48* KO (Extended Data Fig. 3a and b). In contrast, treatment with ATRA, a known inducer of myeloid maturation, induced a stronger differentiation phenotype (Extended Data Fig. 3a and b). Similar results were observed in cell cycle analysis. Cells accumulated in the S-phase after 48h of DAC treatment, but no difference was detected between the *USP48* KO and control cells (Extended Data Fig. 3c).

However, induction of apoptosis as measured by Annexin V/PI staining at 48h and 72h of DAC or GSK-3685032 treatment showed a significant increase in apoptotic/dead cells in the *USP48* KO samples for all three AML cell lines tested (Fig. 1f, Extended Data Fig. 3d). The induction of cell death was confirmed through western blot. Treatment with DAC or GSK-3685032 showed a significant increase in cleaved PARP and cleaved Caspase 3, both markers for apoptotic cell death, in *USP48* KO cells after 48h of drug treatment (Fig. 1g, Extended Data Fig. 3e).

### USP48 alters ubiquitination of H2A variants

To identify direct targets of USP48 and pathways affected by *USP48* loss we utilized a proteogenomic approach. Due to USP48’s role as a deubiquitinating enzyme we first explored changes in ubiquitination, measured by mass spectrometry, as early as three days after induction of *USP48* KO (Extended Data Fig. 4a). Here, we identified histone ubiquitination to be strongly affected by *USP48* KO. Histone 1 linker protein H1.4 (K34/K52) and macroH2A1 (K123/K167) were among the top modified lysine residues in *USP48* KO cells using two different guides (Fig. 2a, Extended Data Fig. 4b). At later time points, we observed an overall increase in histone ubiquitination of H2A variants and H1-type histones (Extended Data Fig. 4c, Extended Data Table 2). While USP48 was previously described as a histone deubiquitinase for H2A (K125/127/129) BRCA1-ubiquitin marks, those sites could not be detected in our assay^18^. MacroH2A1 (K123), however, was also previously identified as a BRCA1 ubiquitin ligase substrate^19^. Furthermore, UIMC1, also known as RAP80, a key component of the BRCA1 complex, was deubiquitinated in *USP48* KO cells (Fig. 2a, Extended Data Fig. 4b and c). The BRCA1-RAP80 complex regulates DNA end resection and is therefore necessary for homologous recombination (HR)^20,21^. In addition to alterations in ubiquitination of histone sites, we observed alterations in ubiquitination in DNA damage repair proteins, particularly PRKDC, XRCC5 (Ku80) and XRCC6 (Ku70), which form the DNA-PK complex (Fig. 2b). DNA-PK recognizes DSBs and is crucial for the choice between the HR and NHEJ DNA damage pathways^22,23^. Cell fractionation experiments detected USP48 in the nuclear and chromatin fraction in the four AML cell lines tested, in line with our finding that *USP48* KO is primarily affecting nuclear proteins (Extended Data Fig. 4d).

**Fig. 2:**
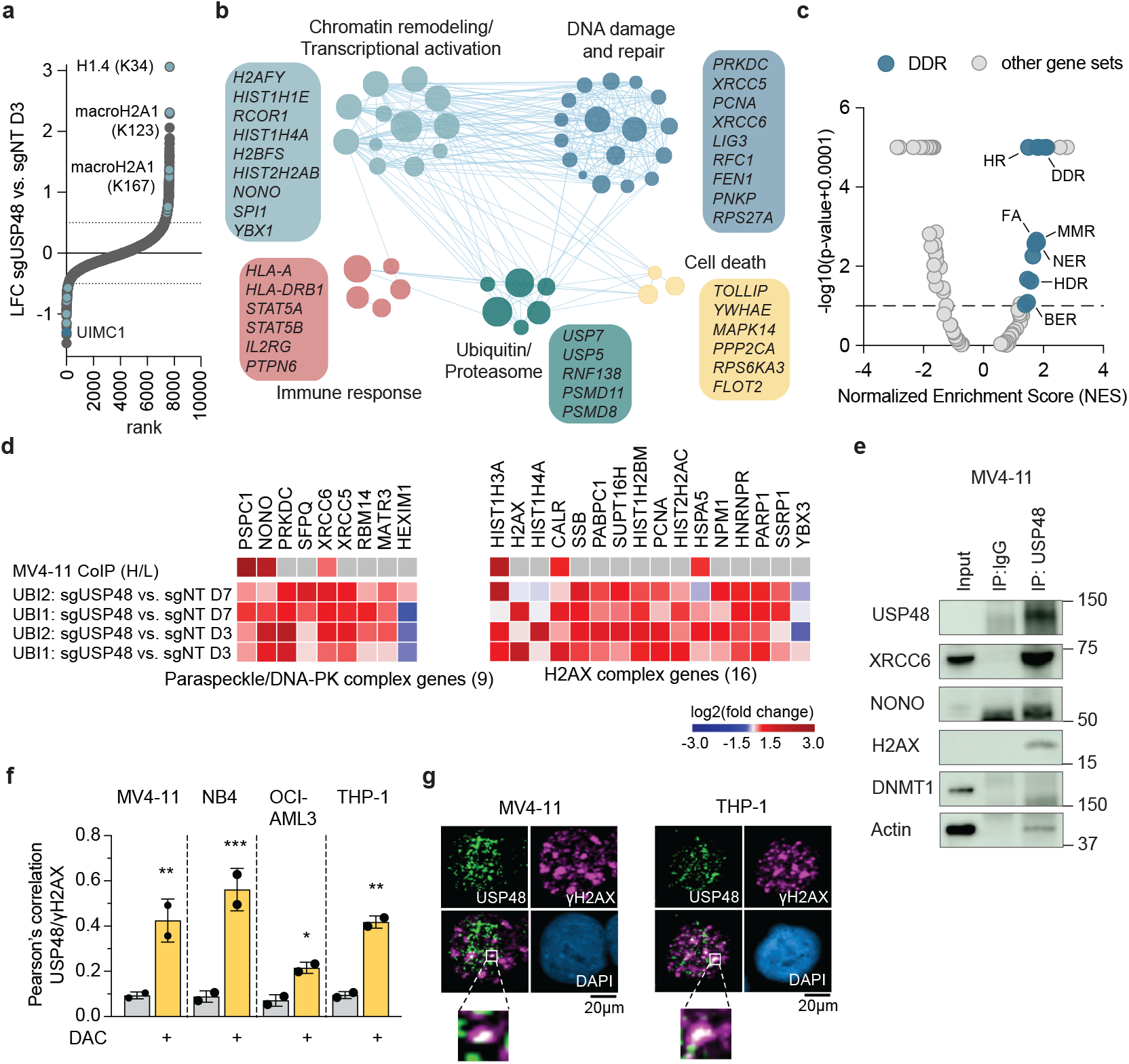
Proteogenomic approach identifies histone and DNA damage proteins as USP48 targets. **a)** Hockey plot of ubiquitinome data in MV4-11 3 days after *USP4*8 KO induction. Each dot represents the log2 fold change (LFC) of one ubiquitin mark identified using mass spectrometry analysis. Significantly increased or decreased histone ubiquitin sites (abs (LFC) ≥ 0.5) are highlighted in light blue. **b)** Enrichment map network of the top functional clusters enriched in the 340 proteins with increased ubiquitination in MV4-11 upon *USP48* KO at day 3. Nodes represent gene sets, with node size indicating the number of genes in each set. Edges connect nodes with a significant overlap (adjusted p-value ≤ 0.05 for the hypergeometric test). Shown in boxes are top leading-edge genes in each functional cluster. Enrichment significance: adjusted p-value ≤ 0.05 for the hypergeometric test. **c)** Volcano plot highlighting the DDR gene sets enriched in genes with increased expression induced by the sgUSP48 vs sgNT at day 3 in RNA-Seq data for MV4-11. Dots represent gene sets in the MSigDB Hallmark collection and in the curated TCGA DDR collection. Significance: abs (NES) ≥ 1.3, p-value ≤ 0.1. **d)** Heatmaps for the paraspeckle/DNA-PK complex (left) and CORUM v3 core histone (H2A/H2B/H3/H4) and H2AX complex (right) enriched in the CoIP, depicting the changes induced in the ubiquitinome UBI1, UBI2 data sets. **e)** Western blot validation of USP48 co-immunoprecipitated proteins in MV4-11 cells in comparison to the input and the IgG control. **f)** Quantifications of USP48-γ2AX colocalization in MV4-11, NB4, OCI-AML3 and THP-1 cells after 48h treatment with 200 nM decitabine (DAC) (n=2), using Pearson’s correlation. 2-way ANOVA multiple comparisons test with Tukey’s corrections was used to compare treatment effect in DAC treated vs. untreated samples * p < 0.05, ** p < 0.01, *** p < 0.001.**g)** Representative IF images of USP48-γ2AX foci in MV4-11 and THP-1 cells upon 48h decitabine treatment (200 nM).

Only moderate changes in transcription and whole proteome analysis were observed with *USP48* KO (Extended Data Fig. 4e and f). These results are consistent with the lack of phenotypic changes observed in AML cells after loss of USP48 alone, where neither cell growth, differentiation, nor viability were affected. The lack of changes in protein abundance at days 3 and 7 after *USP48* loss (Extended Data Fig. 4f, Extended Data Table 3) support the hypothesis that protein ubiquitination through USP48 is functioning as a posttranslational signalling mark and not as a signal for protein degradation. While changes in transcription were mild, gene set enrichment analysis (GSEA) revealed that genes involved in DNA damage repair and chromatin organization were increased in expression, possibly to counteract the effect of USP48 loss (Fig. 2c).

### USP48 co-localizes at DNA damage sites

To further characterize binding partners and targets of USP48, we performed co-immunoprecipitation (CoIP) in MV4-11 cells (Extended Data Table 4). Integrating the ubiquitinome data showed that proteins identified in the CoIP displayed increased ubiquitination upon *USP48* KO in comparison to the full data set, suggesting direct targeting through USP48 (Extended Data Fig. 5a). Complexes annotated by CORUM in the interactome data showed an enrichment of paraspeckle proteins and H2AX complex proteins (Fig. 2d). The core complex members of paraspeckles (NONO and PSPC1) were identified as USP48 binding partners (Fig. 2d). NONO binds to DNA damage sites and plays a role in DNA-PK (XRCC5, XRCC6 and PRKDC) activation; we also observed a change in ubiquitination of each of these DNA-PK complex members with KO of *USP48*^24,25^. The interactions of proteins detected in the mass spectrometry experiment were validated in two AML cell lines using immunoprecipitation followed by western blot. H2AX, NONO and XRCC6 were enriched in the USP48 immunoprecipitated fraction, while no or lower amounts were detected in the IgG control (Fig. 2e, Extended Data Fig. 5b). In comparison, DNMT1 and actin, which are not expected to be bound by USP48, were only present in the input control. To investigate whether USP48 is recruited to DNA damage sites induced by HMAs, we performed immunofluorescence (IF) staining in four AML cell lines treated with DMSO or decitabine (DAC). Treatment with DAC for 48h resulted in an accumulation of γH2AX foci and revealed a significant increase in USP48 foci (Extended Data Fig. 5c and d). Co-localization analysis showed that USP48 overlaps with γH2AX foci at 20–60% of the DNA damage sites as measured by γH2AX (Fig. 2f and g, Extended Data Fig. 5c). This data suggests that USP48 is selectively recruited to a subset of HMA-induced DNA lesions, potentially reflecting damage-site specificity or a differential chromatin context at those sites.

### Combination of HMA treatment with *USP48* KO induces chromatin opening

To better understand USP48’s role in posttranslational histone modification and localization at DNA damage sites, we investigated genome-wide DNA methylation and chromatin accessibility patterns upon *USP48* KO and HMA (GSK-3685032 or DAC) treatment.

Using whole-genome bisulfite sequencing, we found that treatment with low concentrations of GSK-3680532 for 48h led to global demethylation, in both control and *USP48* KO cells, while loss of USP48 alone did not affect DNA methylation (Extended Data Fig. 6a and b). Similarly, no apparent changes in chromatin, accessibility as measured by Assay for Transposase-Accessible Chromatin (ATAC) sequencing, were observed upon *USP48* KO alone (Extended Data Fig. 7a and b). Treatment with GSK-3685032 or DAC in the control cells only had mild effects on opening of chromatin (Extended Data Fig. 7c). In contrast, HMA treatment in *USP48* KO cells showed strong changes in ATAC signal, with focal increases in open chromatin compared to GSK-3685032 or DAC treatment alone (Fig. 3a, Extended Data Fig. 7d). HMA induced ATAC signal changes with *USP48* KO were strongly correlated between the two DNMT1 inhibitors (Extended Data Fig. 7e), with a stronger amplitude in increased signal compared to the *USP48* KO or single agent treatment. Of regions with an increased ATAC signal, 75-85% were identified as *de novo* open chromatin sites (Extended Data Fig. 7f). Conversely, among sites that showed a decrease in ATAC, most sites showed a mild decrease and only 25-33% fell below the detection threshold. Integrating the ATAC signal with DNA methylation profiles revealed that sites showing an increased ATAC signal in the combination of *USP48* KO and GSK-3685032 or DAC were restricted to regions which showed hypermethylation at baseline (Fig. 3b and c).

**Fig. 3:**
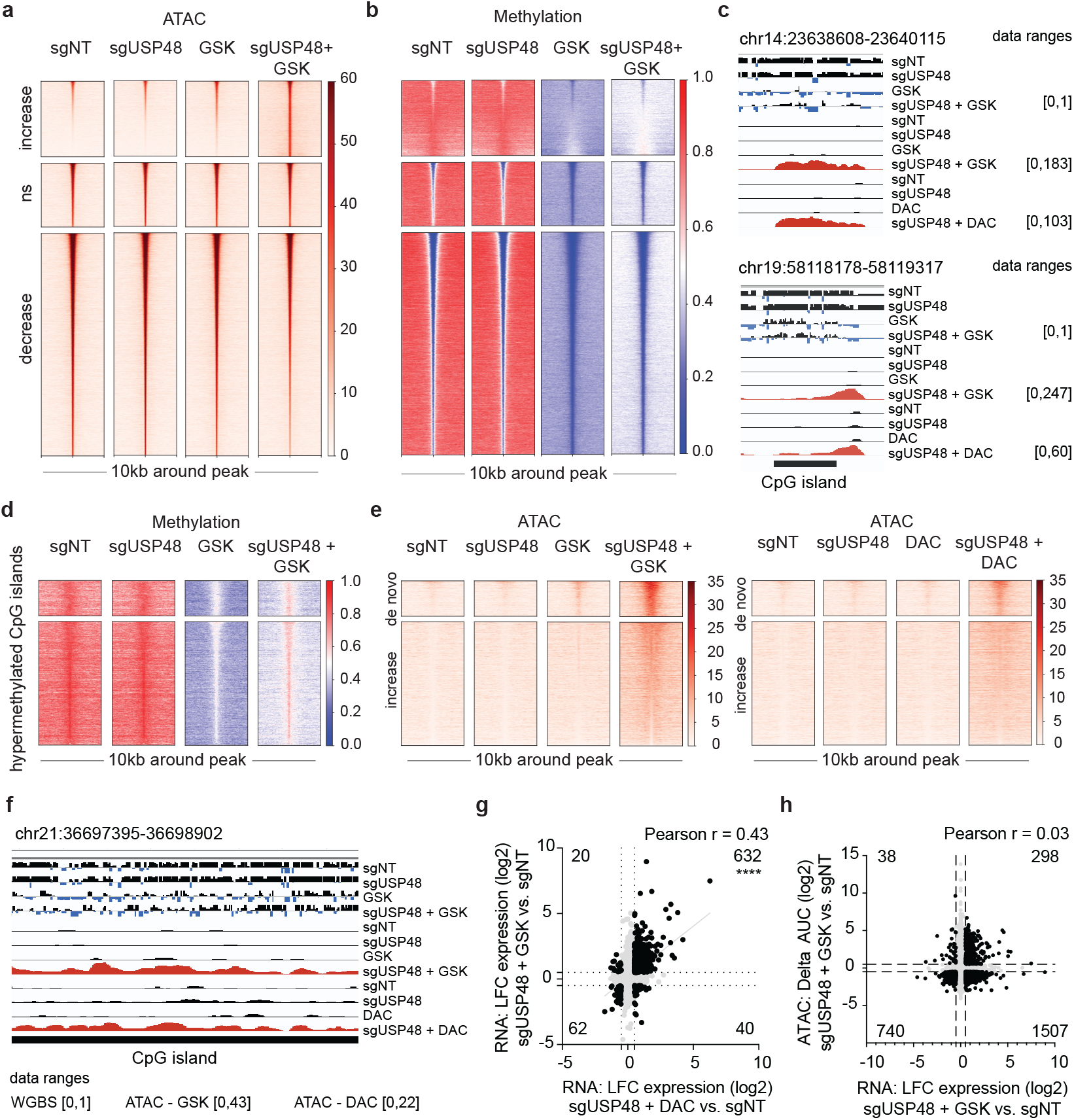
USP48 loss focally increases accessibility of chromatin upon HMA treatment. **a)** Clustered tornado plots for RPKM normalized ATAC signal in regions with increased, not significantly changed, and decreased AUC by sgNT vs. sgUSP48 + GSK-3685032 (GSK). The cutoff for differential binding is set at abs (Δ log2 AUC signal) ≥ 0.5, with a p-value ≤ 0.10. **b)** Tornado plots illustrating the DNA methylation signal in the sgNT vs. sgUSP48 + GSK ATAC-Seq clusters shown in 3a). DNA methylation is represented as halfway normalized beta scores, with scores < 0.5 indicating hypomethylation (blue) and > 0.5 indicating hypermethylation (red). **c)** IGV tracks displaying chromatin accessibility regions from the “ATAC increase” tornado cluster with *de novo* ATAC peak induced by combination treatment in two independent experiments (ATAC - GSK and ATAC - DAC), highlighted in red. Shown are ATAC-seq signal tracks and DNA methylation β-values (midpoint signal) across treatment conditions, with data ranges noted per data set. Co-localized CpG islands are annotated below. **d)** Clustered tornado plots illustrating the DNA methylation signal. Shown are genome-wide CpG island regions that are hypermethylated in the sgNT condition. Top cluster displays hypermethylated CpG islands overlapping with *de novo* ATAC peaks across the treatment conditions. Bottom cluster shows hypermethylated CpG islands with increased ATAC signal below the peak-calling threshold across the treatment conditions. DNA methylation is represented as halfway normalized beta scores, with scores < 0.5 indicating hypomethylation (blue) and scores > 0.5 indicating hypermethylation (red). The CpG islands are ranked by the sgUSP48 + GSK ATAC signal. **e)** Tornado plots illustrating ATAC signal for **left)** sgNT, sgUSP48, GSK and sgUSP48 + GSK treated and **right)** sgNT, sgUSP48, DAC and sgUSP48 + DAC treated MV4-11 on hypermethylated CpG islands shown in 3d). ATAC signal is presented as RPKM normalized, the colour code indicates the signal intensity. Top cluster displays *de novo* ATAC peaks in the sgUSP48 after treatment. Bottom cluster shows ATAC signal at hypermethylated CpG islands with increased ATAC signal below the peak calling threshold. **f)** IGV tracks illustrating CpG islands that are hypermethylated at baseline and show increased chromatin accessibility, below the peak-calling threshold, upon combination treatment in two independent experiments (ATAC - GSK and ATAC - DAC), highlighted in red. Displayed are ATAC-Seq signal tracks and DNA methylation β-values per condition, with data ranges noted per data set and annotated CpG islands highlighted below. **g)** Scatter dot plot illustrating the gene-level correlation between changes in expression induced by sgUSP48 + DAC vs. sgNT at day 4 (x-axis) and sgUSP48 + GSK vs. sgNT at day 2 (y-axis). Dots represent genes. The cutoffs for differential expression are abs (log2 fold change expression) ≥ 0.5, adj p-value ≤ 0.10. Quadrant numbers indicate overlap size. Overlap significance estimated using the two-tailed Fisher exact test **** p < 0.0001. **h)** Scatter dot plot illustrating the correlation between changes in gene expression induced by sgUSP48 + GSK vs. sgNT at day 2 (x-axis) and changes in chromatin accessibility induced by sgUSP48 + GSK vs. sgNT in the ATAC-Seq data at day 2 (y-axis). Each dot represents a gene. The cutoffs for differential expression are: Absolute log2 fold change in expression (abs (log2 fold change expression)) ≥ 0.5. Adjusted p-value ≤ 0.10. The size of overlaps is shown per quadrant.

This genome-wide observation is supported when specifically focusing on the CpG islands across the genome. After separating CpG island that are unmethylated (n=9,590) or fully methylated (n=13,662) in the control cells, only modest changes in chromatin accessibility occur in unmethylated CpG islands after HMA treatment in sgUSP48 cells (Extended Data Fig. 8a and b). In contrast, we observed an accumulation of ATAC signal over CpG islands that are fully methylated at baseline (Fig. 3d-f). A total of 2,021 such regions (14.8%) gained *de novo* ATAC peaks, and an additional 9,590 regions (70.2%) accumulated ATAC signal but below the peak calling threshold (Fig. 3e and f, Extended Data Fig. 8c). The consistent changes observed in sgUSP48 cells with two unique DNMT1 inhibitors suggest that alteration in chromatin accessibility in regions that are highly methylated at baseline is a specific effect of the combination and not an off-target effect of GSK-3685032.

Assessing gene expression changes in *USP48* KO cells upon HMA treatment (GSK-3680532 or DAC) predominantly showed an increased in gene expression, while single agent HMA treatment or *USP48* KO alone showed only mild gene expression changes (Extended Data Fig. 8d and e). As in the case of ATAC signal changes, gene expression changes correlated between DAC and GSK-3685032 treatment in the context of *USP48* KO (Fig. 3f). However, changes in ATAC signal did not correlate with gene expression changes (Fig. 3g), suggesting that the changes in ATAC signal are not concordant with an activation of transcription.

### DNA damage signatures are enriched in USP48 KO combined with HMA treatment

To assess which pathways are activated in *USP48* KO cell upon HMA treatment we analysed transcriptional changes in time course experiments. The mild transcriptional changes previously observed in *USP48* KO cells, with an increase in chromatin and DDR related gene sets, were reproducible in multiple data sets (Fig. 4a, Extended Data Fig. 9a). Low-dose treatment with the two DNMT1 inhibitors led to mild changes in the control cells (sgNT), including the previously described immune response and chromatin changes related to transcriptional activation (Fig. 4a, Extended Data Table 5). In combination with *USP48* KO, the activation of these pathways increased. In contrast to the control cells, the combination of *USP48* KO and GSK-3680532 treatment strongly increased expression of DNA damage repair related genes and induction of cell death pathway signatures (Fig. 4a-c, Extended Data Fig. 9a). The transcriptional programs activated upon GSK-3685032 treatment alone or in combination with *USP48* KO, were reproducible with DAC treatment (Extended Data Fig. 9b-d, Extended Data Table 5). In contrast to GSK-3685032 treatment, single agent DAC induced expression of DDR and cell death related genes in MV4-11 sgNT cells at the later time point, but to a lesser extent than in combination with *USP48* KO cells (Extended Data Fig. 9b). The difference between the two DNMT1 inhibitors could be explained by the DNA-DNMT1 protein adducts that occur with DAC, but not GSK-3685032. In summary, genes related to chromatin remodelling, DDR, immune response, and cell death pathways either increased in expression or were *de-novo* expressed in the combination of DNMT1 inhibitors and *USP48* KO (Fig. 4c, Extended Data Fig. 9d).

**Fig. 4:**
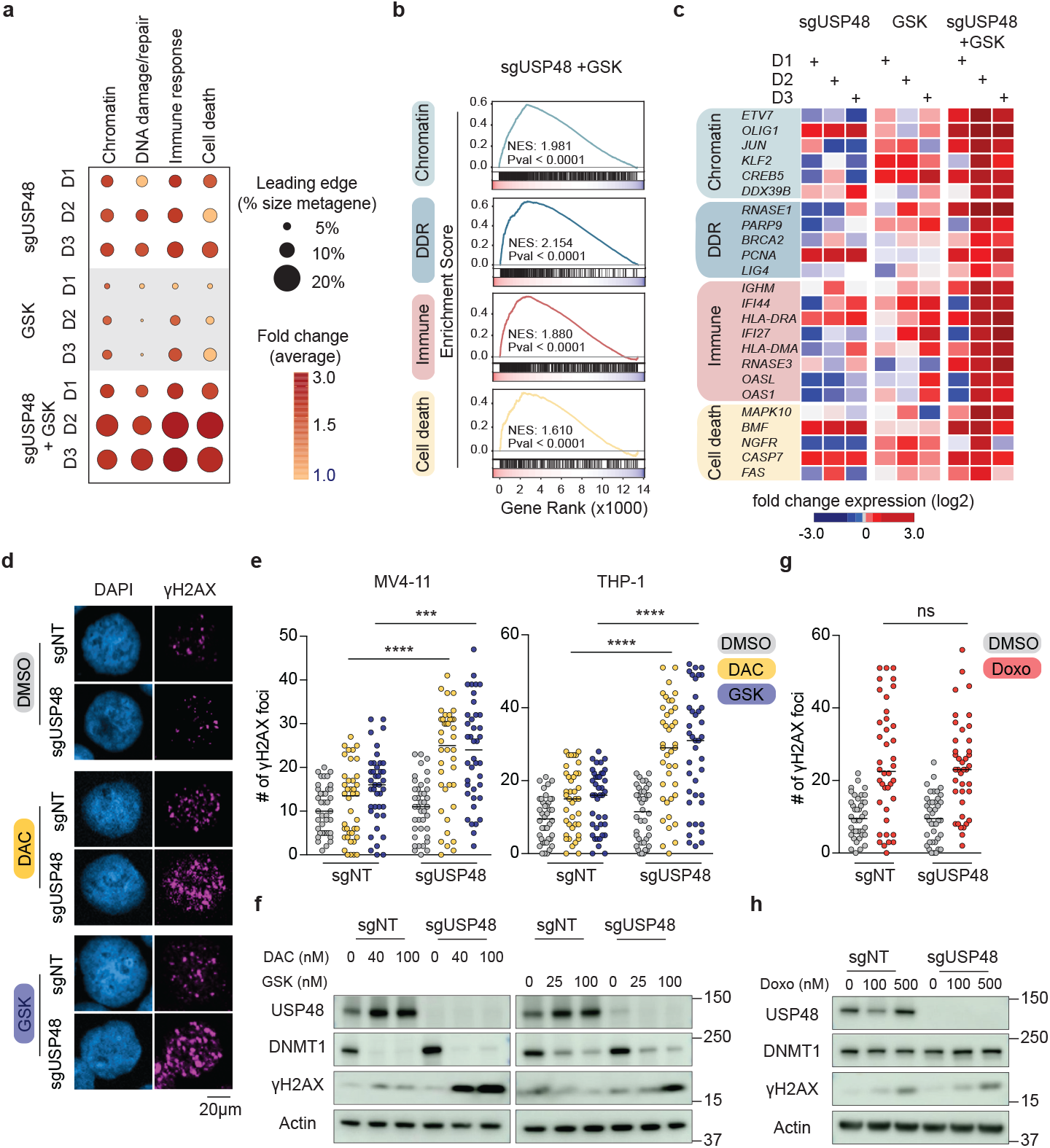
*USP48* KO in combination with HMAs induces DNA damage. **a)** Bubble plot illustrating gene set enrichments associated with genes exhibiting increased expression in sgUSP48, GSK-3680532 (GSK) or sgUSP48 + GSK-3680532 treated conditions. Functional enrichment analysis was conducted using the DAVID platform, encompassing the Gene Ontology and Canonical Pathways (Reactome, KEGG) databases. The plot highlights the top four enriched functional clusters, each represented by a metagene, created by merging the gene sets from the databases that describe the cluster. Bubble size indicates the percentage of genes in the functional cluster metagene with increased expression. Bubble colour reflects the average log2 fold-change expression across the leading-edge genes. **b)** GSEA plots for the 4 metagene sets presented in a) for sgUSP48 + GSK-3680532 at day 2. Significance abs (NES) ≥ 1.3, *P*-value ≤ 0.10, FDR ≤ 0.25. **c)** Heatmap of log2 fold expression changes of top leading-edge genes per functional cluster for sgUSP48, GSK-3680532 treatment alone (GSK) or in sgUSP48 + GSK at day 1, 2 and 3. **d)** Representative images of γH2AX foci in MV4-11 sgNT and sgUSP48 cells after 24h DMSO, decitabine (DAC, 100 nM) or GSK-3680532 (GSK, 25 nM) treatment. DAPI is used as nuclear staining. **e)** Quantification of γH2AX foci in MV4-11 and THP-1 sgNT and sgUSP48 cells upon 24h DMSO (gray), decitabine (MV4-11 100 nM, THP-1 200 nM) or GSK-3680532 (MV4-11 25 nM, THP-1 100 nM) treatment. Each dot represents one nucleus (n = 40 nuclei per sample). 2-way ANOVA multiple comparisons test was used to compare treatment conditions in sgUSP48 vs. sgNT per cell line *** p < 0.001, **** p < 0.0001. **f)** Western blot of MV4-11 sgNT and sgUSP48 cells treated with DAC (40 nM and 100 nM) or GSK-3680532 (25 nM and 100 nM) for 48h. *USP48* KO is validated and levels of DNMT1 and γH2AX are shown. Actin served as a loading control. **g)** Quantification of γH2AX foci in MV4-11 sgNT and sgUSP48 cells upon 6h DMSO (gray) or doxorubicin (500 nM, red) treatment. Each dot represents one nucleus (n = 40 nuclei per sample). 2-way ANOVA multiple comparisons test with Tukey’s correction was used to compare treatment conditions in sgUSP48 vs. sgNT ns – not significant. **h)** Western blot of MV4-11 sgNT and sgUSP48 cells treated with doxorubicin (100 nM and 500 nM) for 6h. *USP48* KO is validated and levels of DNMT1 and γH2AX are shown. Actin served as a loading control.

Following up on the enrichment of DDR signatures in the HMA treated *USP48* KO cells, we assessed the induction of DNA damage, measured by the accumulation of γH2AX using immunofluorescence (IF) staining and western blot. After 24 hours of HMA treatment, the number of γH2AX foci per cell increased significantly in MV4-11 and THP-1 *USP48* KO cells, while only mild changes were observed in the non-targeting controls (Fig. 4d and e). These results were confirmed using western blot analysis, where a dose-dependent increase in γH2AX after 48h of DAC or GSK-3865032 treatment was observed in the *USP48* KO condition in all three AML cell lines (Fig. 4f, Extended Data Fig. 10a). The specificity of γH2AX staining and *USP48* KO for drugs inducing DNA hypomethylation was confirmed using doxorubicin, which induces DNA damage primarily through impairing topoisomerase II (Extended Data Fig. 10b). In contrast to treatment with HMAs, no difference was seen in the number of γH2AX foci per cell and the accumulation of γH2AX in western blot analysis between control and *USP48* KO cells upon doxorubicin treatment (Fig. 4g and h, Extended Data Fig. 10c). These results are in line with the doxorubicin viability data, which showed no differential response between sgNT and sgUSP48 cells (Extended Data Fig. 2d). In addition, western blot analysis showed an increase in USP48 levels upon HMA treatment, but not with doxorubicin treatment (Fig. 4f and h, Extended Data Fig. 10a and c). These results are in line with our finding in IF staining, showing an increase in USP48 foci upon DAC treatment (Extended Data Fig. 5d).

Our results suggest that the accumulation of DNA damage is a secondary effect related to the chromatin changes caused by DNA hypomethylation through HMAs in *USP48* KO cells rather than through conventional DNA damage. The increase in USP48 levels and co-localization to DNA damage sites upon HMA treatment suggests that USP48 is involved in processes counteracting HMA-related structural changes on chromatin and the recognition or repair of the resulting DNA damage.

### *USP48* KO enhances response to HMAs in vivo

To assess USP48 as a potential therapeutic target in AML, we evaluated the effect of *USP48* KO on healthy hematopoietic cell viability and response to HMA treatment. CD34 positive HSPCs were nucleofected with Cas9 protein and guides against *USP48*, a cutting control (Chr2-2), or a positive control (*RPA3*). Colony formation assays and counting experiments showed no significant decrease in cell growth or colony formation upon *USP48* loss in HSPCs from multiple donors (Fig. 5a and b, Extended Data Fig. 11a). In contrast, loss of the essential gene *RPA3* led to growth inhibition (Fig. 5b, Extended Data Fig. 11a). Decreased colony formation was seen in HSPCs treated with DAC, but response rates in control and *USP48* KO cells were similar (Extended Data Fig. 11b and c). In contrast, nucleofected MV4-11 AML cells showed decreased colony formation after 12 days of DAC treatment in combination with *USP48* KO (Extended Data Fig. 11b and c). We confirmed the results using a luminescence readout measuring viability after 96h of treatment with DAC or GSK-3685032 in HSPC sample #2. Again, no difference was observed in response to HMA treatment between sgChr2-2, sgUSP48 #1 and sgUSP48 #2, while sgRPA3 showed the lowest viability (Fig. 5c). These results suggest that loss of USP48 in normal hematopoietic cells does not affect cell growth nor response to HMAs to the same extent as in AML cells.

**Fig. 5:**
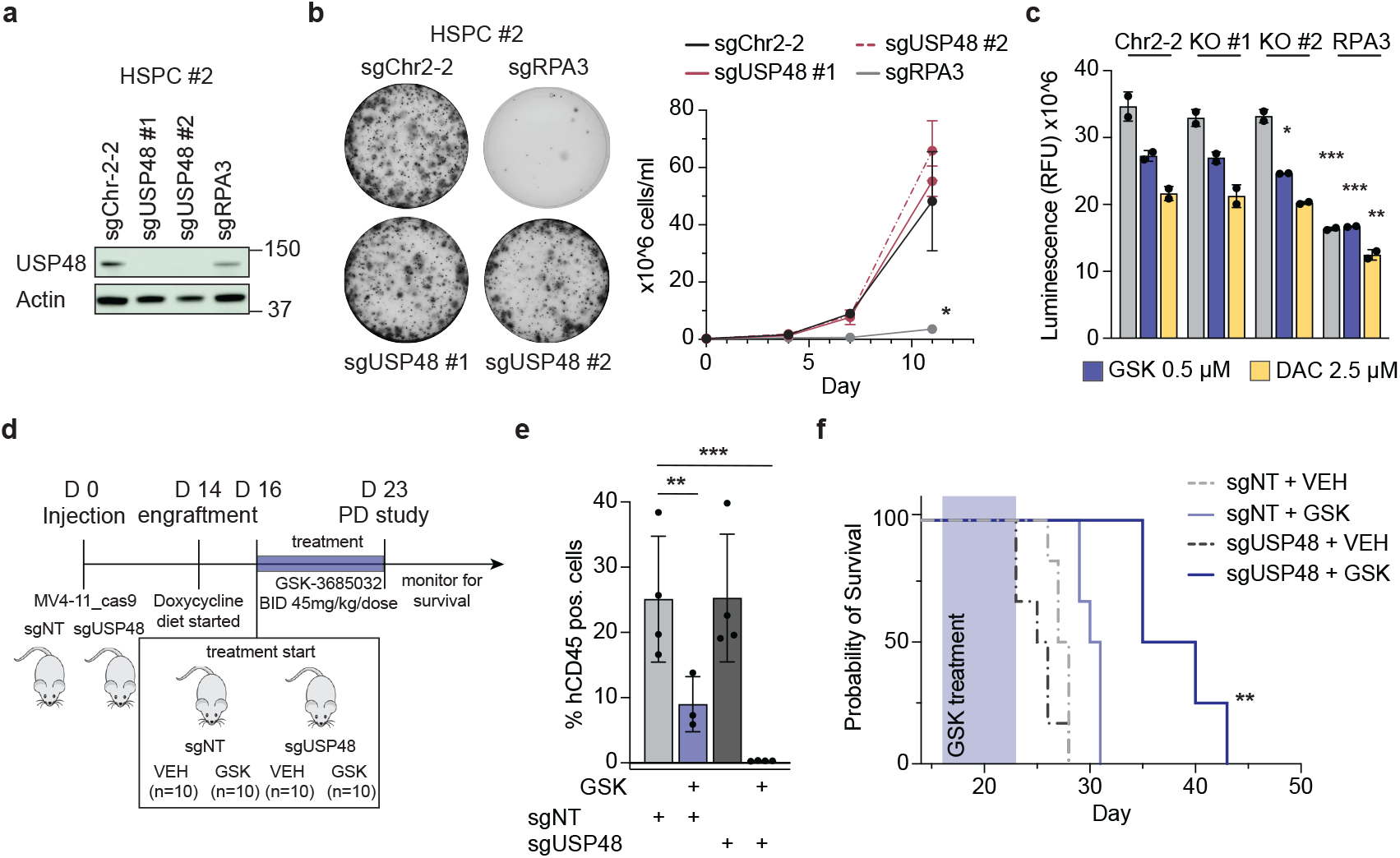
USP48 loss and HMA treatment decrease AML tumour burden *in vivo*. **a)** Validation of *USP48* KO in HSPCs nucleofected with Cas9 protein and sgChr2-2, sgUSP48 #1 and #2 or sgRPA3. Actin served as a loading control. **b)** Representative colony formation assay and counting experiment of HSPC sample #2 carrying sgUSP48 #1 (red), sgUSP48 #2 (red, dashed), sgRPA3 (grey) or sgChr2-2 (black). One-way ANOVA multiple comparisons test with Tukey’s corrections was used to compare KO of *USP48* or *RPA3* vs. sgChr2-2 at Day 11. * p < 0.05. **c)** Cell viability assay in HSPC #2 nucleofected with sgChr2-2, sgUSP48 #1, sgUSP48 #2 or sgRPA3 after 96h of DMSO (grey), GSK-3685032 (blue) or DAC (yellow) treatment. Shown is the CellTiter-Glo luminescence signal per condition (n=2). One-way ANOVA multiple comparisons test with Tukey’s correction was used to compare sgUSP48 or sgRPA3 vs. sgNT per drug condition. * p < 0.05, *\**\* p < 0.01, *** p < 0.001 **d)** Schematic overview of the mouse study performed in NSG mice injected with MV4-11 sgNT or sgUSP48 cells. **e)** Bar graph of % human CD45 positive cells in bone marrow after 7 days of treatment with the vehicle control (VEH, grey) or GSK-3685032 (GSK, blue). Each dot represents the measurement of one mouse. Ordinary one-way ANOVA multiple comparisons test with Tukey’s correction was used to compare treatment conditions vs sgNT + VEH *\**\* p < 0.01, *** p < 0.001. **f)** Kaplan-Meier survival curves of 6 mice per treatment arm (2 mice censored in sgUSP48-GSK group) monitored after 7-day GSK-3685032 treatment. Survival comparison between sgUSP48 + GSK and sgNT + GSK was assessed using the Gehan-Breslow-Wilcoxon test ** p < 0.01.

Next, we determined whether the combination of HMA treatment and *USP48* loss decreases leukaemia burden *in vivo*. Mice were infected with MV4-11 cells carrying guides against *USP48* or a non-targeting control (NT) as well as a fluorescence marker (GFP). After confirmation of engraftment on day 14, a doxycycline diet was initiated for 2 days before GSK-3685032 treatment was started. Mice were then randomized into two arms per group: vehicle (VEH) versus GSK-3685032 (GSK) for a total of 4 groups (Fig. 5d). After 7 days of treatment (45 mg/kg/dose twice per day) 4 mice per group were sacrificed to assess tumour burden. While treatment with the DNMT1 inhibitor had some anti-tumour effect and led to a reduction of AML blasts in the bone marrow, spleen and blood of control mice (sgNT), the combination of USP48 loss with the DNMT1 inhibitor ablated disease burden in all three evaluated tissues (< 0.5% hCD45+ blasts) (Fig. 5e, Extended Data Fig. 12a and b). Two sgUSP48 GSK-3685032-treated mice were censored around one week after treatment end (day 28 and 30) due to dosing-related wounds (scratching at the injection site). Assessment of the bone marrow confirmed the strong anti-tumour effect of the combination, showing <1% of hCD45+ cells in the bone marrow (Extended Data Fig. 12c). The decreased disease burden resulted in prolonged survival for the *USP48* KO + GSK-3685032 group (median survival 37.5 days) in comparison to the vehicle treated mice (*USP48* KO + VEH: 25.5 days, NT + VEH: 27.5 days) or the control mice with GSK-3685032 treatment (30.5 days) (Fig. 5f). Western blot analysis of cells collected prior to doxycycline induction, at day 7 to assess pharmacodynamics, and during monitoring of survival, confirmed *USP48* KO efficiency at day 7 after treatment start, while USP48 protein levels were increased in the *USP48* KO arm at endpoint (Extended Data Fig. 12d). These results suggest that the outgrowth of unedited AML cells in the *USP48* KO group is responsible for the progressive disease and subsequent death of the mice.

## Discussion

Given the importance of HMAs in treating AML/MDS but their modest single agent response rate in patients, our goal in this study was to identify HMA-specific combination targets using CRISPR/Cas9 screening. We tested the two FDA approved HMAs and a novel selective DNMT1 inhibitor (GSK-3685032) and identified an enrichment in HMA sensitizer genes related to chromatin organization, DNA damage repair and transcription, ubiquitination/SUMOylation, and immune response.

The cellular function of the deubiquitinating enzyme USP48, which scored as the strongest sensitizer genome-wide in all five drug treatment conditions in our CRISPR screens, is not well studied but seems to be context specific. Multiple studies describe USP48 as a nuclear located deubiquitinating enzyme involved in cell cycle, NF-κB regulation, and chromatin remodelling in different disease types, leaving the mechanism of *USP48* KO’s combinatorial effect with HMAs in AML in question. Multiple groups independently linked USP48 to BRCA1-specific ubiquitin marks on H2A (K125/127/129), connecting it directly to DSB sites and DNA damage repair pathway choice^18,26–29^. While we could not detect the same histone sites, our data are in line with those findings, showing enriched ubiquitination upon *USP48* KO in a previously identified BRCA1 substrate, macroH2A1 (K123), and co-localization with γH2AX, a marker of DSBs.

To ensure chromatin integrity, cells deploy a fine-tuned network of histone variants and posttranslational modifications (PTMs)^30^. The complex dynamics of different histone variants and PTMs is only partly understood, with signalling networks that seem to be context specific and vary depending on cell cycle phase, chromatin region, and type of DNA damage. Our proteogenomic characterization contributes to the understanding of USP48’s role in the network of epigenetic regulators involved in genome stability and DNA damage and connects USP48 to DNA damage mechanisms directly linked to DNA methylation. Studies have previously shown that DNA demethylation of typically highly compacted chromatin induces chromatin remodelling, which includes increased macroH2A1 incorporation^31^. The non-canonical macroH2A is associated with inactive X chromosomes, but is also found with autosomal chromosomes, correlating with established heterochromatin^32,33^. MacroH2A has a distinct structure; it is almost three times the size of other H2A variants and builds a unique platform for binding partners^34^. We hypothesize that USP48 is crucial in the PTM signalling cascade to ensure transcriptional silencing, chromatin stability, and DNA repair in regions opened upon DNA demethylation. We observed an increase in chromatin accessibility in regions hypermethylated at baseline after *USP48* KO in combination with DAC or GSK3680532 treatment, which was not correlated with transcriptional activation. A possible explanation is that *USP48* KO enhances the previously described effect of HMA treatment in AML cells on chromatin 3D structure, regulatory elements and enhancer hijacking, impacting genome organization and gene expression^35^. Loss of USP48 could also hinder the recruitment of necessary machinery to either 1) stabilize the chromatin or 2) repair DNA lesions caused by the destabilization of the chromatin through DNA demethylation. Both cases would explain the rapid accumulation of DNA damage (γH2AX foci) seen in our experiments. The enrichment of histones and DNA damage related proteins in our ubiquitinome and interactome studies suggests USP48 as an intermediary between macroH2A and DNA damage repair machinery.

Our data suggests that the simultaneous inhibition of DNMT1 and a second target important for heterochromatin organization and histone modifications can enhance the cytotoxic effect of HMAs in AML. Multiple hits in our screen would fit this hypothesis. Among the strongest enriched sensitizers are the histone methyltransferase SETDB1 and its cofactor ATF7IP. Both proteins, as well as TASOR, which also scored among the sensitizers, are part of the HUSH complex and are recruited to transposable elements (TEs) integrated into heterochromatin, hindering their transcription and retrotransposition^36–38^. Silencing of TEs through SETDB1 hinders the uncontrolled expression of TEs, which was previously described to increase chromatin instability and induce an immune response in AML cells^39,40^. Furthermore, multiple members of the histone acetyltransferase (HAT)-module (TADA2B, KAT2A and SGF29) of the SAGA complex score among the HMA sensitizers. The SAGA complex is an important transcriptional regulator and linked to chromatin remodelling during transcriptional activation and at DNA double strand breaks (DSB). Multiple groups have shown that the loss of SAGA modules leads to HR deficiency, and that the complex is required for proper γH2AX formation and DSB repair^41,42^. In addition, studies in yeast described the importance of SAGA subunits SGF29 and SGF73 in the context of establishing heterochromatin boundary formation^43,44^.

To assess USP48 potential in a clinical setting, studies will be needed to understand the effect on normal hematopoietic cells and the synergistic potential with HMAs. In line with a potential therapeutic window, *USP48* is not an essential gene in DepMap, and the KO of *USP48* does not enhance response to HMA in normal HSPCs in our in vitro assays. The universally high expression of *USP48* in AML blasts in multiple patient cohorts (TARGET, Beat AML, TCGA) suggests that USP48 is not a prognostic marker for therapy response.

Ultimately, we will need specific USP48 inhibitors to best evaluate the impact of combining USP48 inhibition with HMAs on AML versus normal tissues. Deubiquitinating enzymes are emerging drug targets. While selective inhibitors for a small number of DUBs, including USP7, USP14, and CSN5 have been reported, the high structural overlap between some DUB family members can pose a challenge to achieving selectivity^45–47^. Various groups reported compounds with USP48 binding affinity ex vivo at high molarity or non-selective inhibition; optimization of these molecules is needed for *in vitro* and *in vivo* applicability^48,49^. Efforts to identify potent, specific USP48 inhibitors are underway.

In summary, we identified USP48 as a therapeutic target for combination with HMAs. Our data confirm USP48 as a histone deubiquitinase, influencing the modification of H2A variants and other proteins important for chromatin structure and DNA damage repair pathways. While *USP48* KO alone does not affect cell viability, combining *USP48* KO with HMA treatment is highly lethal to AML, suggesting a synergistic interplay between DNA hypomethylation, posttranslational ubiquitin modifications, and DNA damage in the setting of AML biology. USP48 is thus a new target for combination therapy with HMAs for AML.

## Acknowledgements

C.S. was supported by a grant from the German Research Foundation (Deutsche Forschungsgemeinschaft [DFG], (SCHN 1622/1-1)) and by a Helen Gurley Brown Fellowship. C. S. and K. S. were supported with funding from The Selig Family Fund for Pediatric Cancer Research. R.B.S. and R.E.G. were funded by a Hyundai Hope on Wheels HOPE grant for Childhood Cancer Research. Y.R. was supported by the HHMI Hanna H. Gray Fellowship during the duration of this study. K.S. was supported by a National Cancer Institute R35 CA283977 and the Children’s Leukemia Research Association. Fig. 1a was created with BioRender. The results published here are in part based upon data generated by the Cancer Dependency Map and TCGA Research Network. We thank Sarah Buhrlage for discussions about DUB inhibitors.

## Author contributions

Conceptualization: C.S., K.S.; Methodology: C.S., M.L.S., R.B.S., A.T.B., Y.R., B.H.; Experiments: C.S., L.A.M., A.M., M.L.S., R.B.S., A.T.B., A.T. S.S., B.H.; Computational analysis and data interpretation: G.A., C.S., F.B., S.L., V.H., K.S.; Visualization: C.S., G.A., M.L.S.; Funding acquisition: K.S. and C.S.; Resources: R.E.G., D.E.R., T.O., D.C.; Supervision: K.S., C.S.; Writing C.S, K.S and L.M., with input from all authors.

## Materials and methods

### In vitro culture of cell lines

MV4-11 (CVCL_0064) cells were generously provided by Scott Armstrong at Dana-Farber Cancer Institute. OCI-AML3 (CVCL_1844) and NB4 (CVCL_0005) were purchased from DSMZ, and THP-1 (CVCL_0006) was purchased from ATCC. All AML cell lines were cultured in RPMI medium (Gibco) supplemented with 10 % FBS (Sigma-Aldrich) and 1 % penicillin-streptomycin (P/S) (Life Technologies). Cell lines were cultured for up to three months following thawing for experimental use. CD34^+^ hematopoietic stem/progenitor cells (HSPC) (Lonza) were briefly cultured in StemSpan II media (STEMCELL Technologies) supplemented with 100 ng/ml human SCF, TPO, and FLT3L, and 10 ng/ml IL3 and IL6. The identities of parental cell lines and newly generated models were validated by short tandem repeat profiling at the Molecular Diagnostics Laboratory at the Dana-Farber Cancer Institute and tested for Mycoplasma contamination using the MycoAlert Mycoplasma Detection Kit (Lonza, LT07-318).

### CRISPR/Cas9 screen

MV4-11 Cas9-mCherry positive cells were transduced with the Avana CRISPR knockout library (lentiCRISPRv2) in two biological replicates to achieve 30-40 % infection efficiency. Twenty-four hours after transduction, cells were pooled and selected with puromycin (Gibco) (2 µg/ml) for 72 hours. After selection, cells were counted and split into three sets of 50×10^6^ cells per replicate to maintain a library representation of >500 cells per sgRNA. Treatment with DMSO (Thermo Fisher), 200 nM azacitidine (Sigma Aldrich, A3656), or 100 nM decitabine (Tocris Bioscience, 26-241-0) started on day 7 after transduction. Cells were counted, split, and retreated every 4 days; surviving cells were harvested after 16 days. For additional screening using lower concentrations (IC10) cells were treated as described above. After puro selection cells were split into four treatment conditions per replicate: DMSO, 100 nM azacitidine, 40 nM decitabine, or 25 nM GSK-3685032 (MedChem Express, 50-226-0838). Genomic DNA was extracted from collected cell pellets using a NucleoSpin Blood L kit (Takara #740954.20). sgRNA sequences from the AVANA-4 library were PCR-amplified and subjected to standard Illumina sequencing at the Genomic Perturbation Platform (Broad Institute). sgRNA read counts were deconvoluted using the PoolQ software (https://portals.broadinstitute.org/gpp/public/software/poolq).

Annotation files for sgRNA efficacy (CRISPRInferredGuideEfficacy.csv) and common essential genes (CRISPRInferredCommonEssentials.csv, AchillesCommonEssentialControls.csv) were retrieved from the 24Q2 DepMap public database (https://depmap.org/portal/download/). Raw sgRNA counts were log-normalized to the initial plasmid DNA (pDNA) reference. Guides with low efficacy (≤ 0.30) or poor representation in pDNA were excluded. The remaining sgRNAs were mapped to genes using the CP0033_GRCh38_NCBI_CRISPRko_strict_gene_20221209.chip annotation. Nonessential genes served as negative controls, while common essential genes were positive controls for quality assessment. Replicate reliability was evaluated via scatter plots; all samples passed QC. *Differential Dependency Analysis*. Differential gene dependency scores were computed for hypomethylating agent (HMA)-treated conditions relative to DMSO controls, using log-normalized sgRNA data averaged across replicates. A hypergeometric distribution-based method (https://portals.broadinstitute.org/gpp/public) was applied to assess enrichment of gene-level effects. Gene-level scores represented the average log-normalized dependency of corresponding sgRNAs. P-values were calculated from the hypergeometric probability mass function of ranked dependencies. Genes meeting thresholds of |log-fold change| ≥ 0.5 and p-value ≤ 0.10 were considered significant. Genes with negative differential scores in drug vs. DMSO conditions were categorized as sensitizers (KO enhances drug sensitivity), whereas those with positive scores were associated with drug resistance (KO reduces drug sensitivity).

#### Consensus Sensitizer Set and Enrichment Mapping

A consensus common sensitizer list of 300 genes was created based on the genes that scored as sensitizers for at least two of the three HMA drugs in either IC30 or IC10 treated samples. Functional enrichment clustering for Gene Ontology terms and canonical pathways (Reactome, KEGG) in the sensitizer consensus list was performed using the DAVID platform^1^. Significantly enriched gene sets (adjusted p-value ≤ 0.05 for the hypergeometric test) were visualized using the Enrichment Map^2^ available from the Cytoscape platform^3^. Nodes grouped in functional clusters represent enriched gene sets, with size proportional to gene count; edges connecting nodes indicate significant node gene set overlap (p ≤ 0.05 for the hypergeometric test).

### Cloning and lentiviral transduction

Cells with constitutive and stable Cas9 expression were created. To generate CRISPR/Cas9 knockouts, single-guide RNAs (sgRNA) targeting genes of interest were cloned into a vector containing doxycycline inducible sgRNA and constitutive GFP for an inducible system (Addgene plasmid #70183). The sgRNA sequences that were used are: non-targeting (NT) (CCGCGCATTTCAGAGCACAA), sgUSP48#1 (TTTGTGGGCCTGACTAACCT), sgUSP48 #2 (TCGATGATCCCAACTGTGAG). Lentiviral particles were produced using TransIT (Mirus) in HEK293T cells. Virus was harvested 48 hours following transfection, and cells were transduced using lentivirus and polybrene (Santa Cruz Biotechnology) (8 µg/mL) through a spin infection (2,000 rpm at 30 °C for 1.5 hours).

### Cell viability assay

Cells carrying Cas9 and sgRNA GFP-inducible guides were doxycycline induced (200 ng/ml) and incubated at 37 °C for three days. Induced cells were then plated in 384-well plates with 50 µl per well at a density of 0.2×10^6^ cells/ml. A drug dispenser (D300e Digital Dispenser, HP) was used to add serial drug dilutions before plates were incubated for 96 hours at 37 °C. At the time of read-out, 10 µl CellTiter-Glo (Promega) was added to each well and plates, protected from light, were incubated for 15 minutes before luminescence was read on either a FLUOstar OPTIMA (BMG Labtech) or a CLARIOstar^*Plus*^ (BMG Labtech) plate reader.

### Annexin V/PI staining

Cells carrying Cas9 and sgRNA GFP-inducible guides were induced with doxycycline (200 ng/ml) and incubated at 37 °C for three days. Induced cells were then plated in a 12-well plate at a density of 0.3×10^6^ cells/ml in triplicates, treated with drug, and incubated at 37 °C for 48 or 72 hours. Following incubation, cells were stained using the BioLegend APC Annexin V apoptosis detection kit with PI (propidium iodide). Following two brief washes in PBS + 1 % FBS, cells were resuspended in 100 µl Annexin V Binding Buffer with 10 µl PI solution and 5 µl APC-conjugated Annexin V, vortexed, and incubated for 15 minutes at room temperature protected from light. Cells were then topped off with 400 µl Annexin V Binding Buffer and analysed by flow cytometry (BD FACSCelesta). Data were analysed using FloJo.

### Immunoblotting

Cells were lysed using RIPA buffer (Thermo Fisher) supplemented with a complete Protease Inhibitor Cocktail tablet (Roche) and PhosSTOP (Roche) tablet on ice. The lysates were quantified using a bicinchoninic acid (BCA) assay (Thermo Fisher), normalized, and diluted with RIPA and 4x Laemmli buffer (Bio-Rad Laboratories) supplemented with β-mercaptoethanol (Sigma-Aldrich). Proteins were then separated using 4-12% SDS-PAGE gels (Invitrogen) and transferred to methanol-activated polyvinylidene difluoride membranes (Millipore). Following the transfer, unspecific binding was blocked using 5 % milk (Cell Signaling Technologies) in TBST. Primary antibodies were diluted in TBST + 5% BSA and incubated with membranes overnight at 4 °C. The following antibodies were used: USP48 (A301-190A, Bethyl, 12076-1-AP, Proteintech), actin (4970S, Cell Signaling Technologies), DNMT1 (PA1-880, Invitrogen), γH2AX (2577S, Cell Signaling Technologies), Caspase3 (9662S, Cell Signaling Technologies), cCaspase3 (9664S, Cell Signaling Technologies), PARP (9542S, Cell Signaling Technologies; 436400, Invitrogen), XRCC6/Ku70 (10723-1-AP, Proteintech), XRCC5/Ku80 (16389-1-AP, Proteintech), HDAC1 (2062S, Cell Signaling Technologies), H3K27me3 (9733S, Cell Signaling Technologies), H2AX (7631S, Cell Signaling Technologies), NONO (11058-1-AP, Proteintech). Membranes were washed in TBST following incubation with primary antibodies and incubated with HRP-linked goat anti-rabbit IgG and anti-mouse IgG (CST #7074, #7076) secondary antibodies, diluted in 5 % milk, before being imaged using Super Signal West Femto Maximum Sensitivity Substrate (Thermo Fisher Scientific) on an Amersham Imager 680.

### Differentiation assays

For morphology staining, doxycycline induced cells were plated at a density of 0.3×10^6^ cells/ml, treated and incubated for 72 hours at 37 °C. 50,000 cells were harvested and resuspended in 100 µl PBS (Gibco). Cells were spun onto slides (5 minutes at 900 rpm) using a

Shandon Cytospin 4 (Thermo Electron Corporation) and left to dry for 30 minutes at room temperature. Slides were then fixed in methanol for 5 minutes before incubation in May-Grünwald stain (Sigma-Aldrich) for 5 minutes. Slides were washed with PBS (5 minutes) and incubated in 5% Giemsa stain (Sigma-Aldrich) for 20 minutes before being rinsed with water. Coverslips were mounted using one drop of mounting media (Shandon-mount, Epredia), and slides were imaged using a digital microscope (Olympus BX41).

For assessment of CD11b expression, cells were plated at a density of 0.3×10^6^ cells/ml and treated for 72 hours. 0.4×10^6^ cells were harvested washed and stained with CD11b-APC (BioLegend, #101212) antibody (1:100 in flow buffer (PBS+ 2 % FBS)) for 20 minutes at room temperature. Cells were washed once in flow buffer, resuspended, measured by flow cytometry (BD FACSCelesta) and analysed using FlowJo.

### Cell cycle

Doxycycline induced cells were seeded at a density of 0.3×10^6^ cells/ml in 12 well plates, treated with HMAs, and incubated at 37 °C for 48h. At the desired timepoint, cells were labelled for 90 minutes with 10 µmol/L EdU at 37 °C, harvested, and prepared using the Click-iT EdU Alexa

Fluor 647 kit (C10636, Life Technologies). After incubating samples with FxCycle Violet DNA staining (F10347, Thermo Fisher Scientific) for 20 minutes at room temperature, samples were measured using flow cytometry (BD FACSCelesta) and data were analysed using FlowJo.

### CoIP/Interactome

50×10^6^ cells were harvested per condition, spun down, and resuspended in 37.5 ml of FBS free medium. 2.5 ml of 16 % formaldehyde (Thermo Fisher Scientific) was added to cells, for a final concentration of 1 % formaldehyde, and incubated for 7 minutes at room temperature. 4 ml of 2.5 M glycine was added to the mixture to halt the crosslinking reaction. Cells were spun down for 5 minutes at 500g, resuspended in 1ml of modified RIPA lysis buffer, and incubated on ice for 10 minutes. Following incubation, cells were spun down for 10 minutes at 15000 rpm at 4 °C, and the lysate was transferred to a fresh tube. Protein concentration was measured using a BCA assay, and samples were diluted to a concentration of 2 mg/ml. 60 µl of each sample was removed and kept as an input control. Per cell line, samples were split and 10 µg IgG (2729S, Cell Signaling) or USP48 (A301-190A, Bethyl) antibody was added to the samples, and incubated overnight rotating at 4 °C. The following morning, 40 µl Protein A/G Plus Agarose beads for Immunoprecipitation (Thermo Fisher Pierce) were added to samples and incubated rotating at 4 °C for 4-6 hours. Cells were then spun down at 500g for 5 minutes at 4 °C. Beads were washed three times with 500 µl lysis buffer, incubating samples on ice for 5 minutes with each wash step before spinning for 5 minutes at 500g at 4 °C. Following washing, beads were boiled in 25 µl 4x LDS buffer (NuPAGE) and 5 µl 10x reducing agent (NuPAGE) for 5 minutes at 95°C, the supernatant collected and analysed by immunoblotting.

For mass spectrometry analysis cells were grown in SILAC medium for 14 days. Preparation of protein pull-down samples was done as described above. “Light” (IgG) and “Heavy” (USP48) labelled eluates were pooled 1:1 and subjected to SDS-PAGE. The lanes of separated proteins were stained with Coomassie Brilliant Blue and excised as 23 individual gel pieces. After reduction with 10 mM DTT and alkylation with 55 mM iodoacetamide in the gel matrix the protein samples were subjected to in-gel digestion with trypsin (Promega) overnight. The peptides were extracted and analysed by LC/MS as described below.

### Proteome/Ubiquitinome

For whole proteome and ubiquitinome analysis cells were plated at a density of 0.3×10^6^ cells/ml and treated with doxycycline. Proteins were collected at day 3 and day 7 after induction of knockout. Per condition 100×10^6^ cells were collected, lysed by suspension in urea buffer (8 M urea, 20 mM HEPES pH 8.0, 1 mM sodium orthovanadate, 2.5 mM sodium pyrophosphate, 1 mM beta-glycerophosphate) and 3 sonication cycles. Protein concentration in the cleared lysate was determined using Pierce 660 nm assay (Thermo Fisher Scientific) and the samples were reduced with DTT for 1 hour at 37 °C (final concentration 5 mM) and alkylated with iodoacetamide for 30 min at room temperature and protected from light (final concentration 10 mM). After dilution to 1 M urea with 20 mM HEPES pH 8.0 the samples were digested overnight with trypsin in an enzyme-to-substrate ratio of 1:100, followed by peptide purification using C18 cartridges (Sep-Pak classic 360 mg, Waters). Peptide concentration in the eluates was determined using Pierce Fluorometric Assay (Thermo Fisher Scientific) and 10 µg or 3 mg per condition were taken for global proteome profiling or enrichment of ubiquitinated peptides, respectively. For the latter, the samples were processed using the PTMScan HS Ubiquitin remnant kit (Cell Signaling) that utilizes the diglycyl lysine antibody coupled to magnetic beads. Both the proteome and ubiquitinome samples were dried in a vacuum centrifuge, dissolved in 20 mM HEPES pH 8.0 and labelled for 1h at room temperature with 100 µg of TMT10plex reagents (Thermo Fisher Scientific). After quenching the labelling reactions for 15 min with hydroxylamine in a final concentration of 0.2 %, the individually labelled samples were combined. Both multiplexes were dried by vacuum centrifugation and separated into 8 fractions using Pierce High pH RP fractionation kit (Thermo Fisher Scientific). The peptide samples were vacuum dried and for LC/MS analysis dissolved in 0.1 % FA.

### LC/MS analysis and data processing

The peptide samples were analysed on a nano-UHPLC system (Ultimate 3000 nRSLC, Thermo Fisher Scientific) coupled to a quadrupole-Orbitrap hybrid mass spectrometer (Q Exactive HF, Thermo Fisher Scientific) via a nano-ESI ion source (Nanospray Flex, Thermo Fisher Scientific). After clean-up on a C18 pre-column (5 cm, packed with ReproSil-Pur 120 C18-AQ, 5 µm particle size, Dr. Maisch GmbH), the samples were separated on a C18 analytical column (32 cm, packed with ReproSil-Pur 120 C18-AQ, 1.9 µm particle size, Dr. Maisch GmbH) with a linear gradient of 2 to 40 % solvent B (80 % ACN, 0.1 % FA) against solvent A (0.1 % FA) over 118 min at a flow rate of 300 nl/min. The peptides eluting from the column were ionized and analysed by tandem MS (MS/MS) in a data-dependent acquisition scheme. Full scans of the ion population were acquired with the Orbitrap analyser in the range of m/z 350-1600 with a resolution setting of 120,000 FWHM at m/z 200 and considering charge states 2-6. The 20 most abundant precursor ions were selected for collision-induced dissociation (HCD) with a normalized collision energy of 32 % (isolation window m/z 1.4). Fragment ion spectra were recorded using the Orbitrap with a resolution setting of 15,000 FWHM at m/z 200 (SILAC) or with a fixed first mass of m/z 110 and a resolution setting of 60,000 FWHM at m/z 200 (TMT). AGC target values and maximum injection times for MS and MS/MS were set to 1×10^6^ in 40 ms and 1×10^5^ in 64 ms, respectively. Fragmented precursor ions were excluded from repeated isolation for 20 s.

For protein identification and quantification, the raw data was processed with version 1.6.17.0 of the MaxQuant software^4^. The mass spectra were matched against the Uniprot human protein database (Swiss-Prot, downloaded 02-2021) and a collection of frequently observed laboratory contaminants using the integrated Andromeda peptide search engine. The mass tolerances were set to 4.5 ppm and 20 ppm for precursor and fragment ions, respectively. Acetylation of the protein N-terminus and oxidation of methionine were considered as variable modifications and cysteine carbamidomethylation was set as fixed modification. For ubiquitinome analysis the diglycyl remnant on lysine was defined as an additional variable modification. The minimal peptide length was set to seven amino acids and to up to two missed tryptic cleavage sites were allowed. Both peptide and protein FDR were set to 0.01 using a decoy approach by searching the reversed database. For SILAC-based quantification the label multiplicity was set to value 2 addressing the light (Arg-0/Lys-0) and heavy (Arg-10/Lys-8) conditions. The initial MaxQuant output data was filtered by removing potential contaminants, hits to the decoy database and proteins identified solely based on modified peptides. The TMT quantitation data was adjusted for equal peptide loading within multiplexes by standardization on the median summed-up reporter intensities of each TMT channel. Replicate injections were scaled using factors calculated from a mock reference that was obtained from the median reporter intensities of each protein.

### RNA sequencing

Cell pellets were collected following treatment, and RNA was isolated using the RNeasy Plus Mini Kit (Qiagen). Cells were harvested and purified using on column gDNA elimination. RNA concentration was determined using a NanoDrop (Thermo Fisher Scientific). RNA samples were sent for mRNA library preparation and mRNA sequencing as 150 bp PE reads at Novogene.

### Immunofluorescence staining

Cells were seeded at a density of 0.2×10^6^ cells/ml. The following day, cells were treated with HMAs as desired and incubated for 24-48 hours at 37°C. Doxorubicin was incubated for 6 hours at a concentration of 500 nM before processing of the cells. Cells were harvested, washed once with PBS, and resuspended in 200 µl PBS. 100 µl of the cell mixture was spun onto slides (5 minutes at 400 rpm) using a Shandon Cytospin 4 (Thermo Electron Corporation). Cells were pre-extracted by incubating in CSK buffer (10 mM HEPES, 100 mM NaCl, 300 mM sucrose, 3 mM MgCl_2_) plus 0.3 % Triton-X100 on ice for 10 minutes. Following pre-extraction, cells were washed once in PBS and fixed by incubating in 4 % paraformaldehyde for 20 minutes at RT. Cells were then rinsed twice with PBS and permeabilized and blocked by incubating in 3 % BSA in PBS plus 0.5 % Triton-X100 for 30 minutes at RT. Slides were incubated in primary antibody, diluted in 3 % BSA in PBS (USP48: A301-190A, Bethy, 1:750, γH2AX: 05-636, Millipore, 1:3000), overnight at 4 °C under humidity. The following day, slides were washed with PBS three times, 5 minute incubation per wash, and incubated in secondary antibody diluted in 3% BSA in PBS (A10042, Invitrogen, Donkey anti-Rabbit IgG (H+L) Highly Cross-Adsorbed Secondary Antibody, Alexa Fluor 568, 1:1000, A31571, Invitrogen, Donkey anti-Mouse IgG (H+L) Highly Cross-Adsorbed Secondary Antibody, Alexa Fluor 647, 1:1000) for one hour at RT protected from light. Following incubation, slides were washed three times in PBS, incubating for 5 minutes with each wash, and coverslips were mounted using DAPI containing mounting media (Abcam). Slides were left to dry overnight at room temperature and then stored at 4 °C. Images were acquired with a Zeiss Axio Observer microscope equipped with a digital camera at 63x magnification. Immunofluorescence foci were quantified in ImageJ (v.1.53a). All statistical analyses were performed by Prism v9.5 (GraphPad).

### Cell fractionation

2×10^6^ cells per sample were collected, washed in PBS and cell fractionation was performed using the Subcellular Protein Fractionation Kit for Cultured Cells (Thermo Fisher Scientific). The cytoplasmatic, soluble nuclear and chromatin bound fractions were collected und transferred into fresh tubes. Proteins were processed for immunoblotting as previously described.

### Whole-genome bisulfite sequencing (WGBS)

Doxycycline induced cells were treated with 100 nM GSK-3685032 for 48 hours. Following treatment, cells were harvested, and DNA was extracted using a DNeasy Blood and Tissue Kit (Qiagen). Samples were then sent to Novogene for QC and sequencing. First, genomic DNA spiked with lambda DNA were fragmented to 200-400 bp, bisulfite treated, and methylation sequencing adapters were ligated, followed by double strand DNA synthesis. The library was ready after size selection and PCR amplification. The library was checked with Qubit and real-time PCR for quantification and bioanalyzer for size distribution detection. Quantified libraries were pooled, and a total of 8 libraries were Illumina sequenced as 150 bp PE reads, aiming for 10-fold genomic coverage (∼200 million PE reads).

### ATAC seq

Sample preparation for ATAC-sequencing was carried out using an Active Motif kit. 50,000 cells per condition were washed once in ice cold PBS and resuspended in 100 µl of ice cold ATAC lysis buffer, transferred to a PCR tube and spun down. Nuclei were resuspended in 50 µl Tagmentation Master Mix (25 µl 2x Tagmentation buffer, 2 µl 10x PBS, 0.5 µl 1.0% Digitonin, 0.5 µl 10% Tween 20, 12 µl H_2_O, 10 µl assembled transposomes) and incubated for 30 minutes in a thermomixer set to 37 °C and 800 rpm. Following incubation, samples were transferred to fresh PCR tubes, and 250 µl DNA Purification Binding Buffer and 5 µl 3M sodium acetate were added to each sample. DNA was then purified by transferring samples to DNA purification columns, washing with Wash Buffer, and eluting in DNA Purification Elution Buffer. Each sample was assigned a unique primer pair to be used in the PCR amplification of Tagmented DNA (33.5 µl Tagmented DNA, 2.5 µl of 25 µM i7 indexed primer, 2.5 µl of 25 µM i5 indexed primer, 1 µl of 10 mM dNTPs, 10 µl 5X Q5 reaction buffer, 0.5 µl of 2U/µL Q5 polymerase; 72 °C for 5 minutes, 98 °C for 30 seconds, 10 cycles of 98 °C for 10 seconds, 63 °C for 30 seconds, and 72 °C for 1 minute). PCR products were cleaned using SPRI bead clean-up, and samples were sequenced as 2PE 150 bp reads on an Illumina NovaSeq by the Genomics core facility at the Dana-Farber Cancer Institute.

### Nucleofection

Nucleofection of CD34+ HSPCs was performed using the P3 Primary Cell 4D-Nucleofector X Kit S (Lonza). CD34+ HSPCs (Lonza, 2M-101C) were thawed and stimulated in StemSpan II media supplemented with 100 ng/ml human SCF, TPO and FLT3L, and 10 ng/ml IL-3 and IL-6 for two days prior to the experiment. Per transfection, 6 µg purified Cas9 nuclease (IDT) was mixed with 100 pmol chemically modified synthetic sgRNA (Synthego, sgRNA target sequence: sgChr2-2 (GGTGTGCGTATGAAGCAGTG), sgRPA3 (GATGAATTGAGCTAGCATGC), sgUSP48#1 (TTTGTGGGCCTGACTAACCT), sgUSP48 #2 (TCGATGATCCCAACTGTGAG)) in P3 buffer and incubated for 10 minutes at RT to form RNP. Cells were counted, washed, resuspended in P3 buffer (0.25×10^6^ cells per reaction), and transferred to the RNP mixture. 25 µl of the cell + RNP mix was pipetted into one well of a 16-well Nucleocuvette Strip and electroporated on the 4D-Nucleofector (Lonza) with the program DZ-100. The nucleofected cells were then transferred into prewarmed media in a 24-well plate and incubated at 37 °C for two days.

### Colony formation

Nucleofected cells were counted and resuspended in MethoCult Express methylcellulose media (StemCell Technologies #04437) at a density of 2000-3000 cells/ml. Resuspended cells were vortexed and left at RT to allow the bubbles to dissipate. In triplicates, 1 ml of the cell/methylcellulose mix was then plated onto 35 mm cell plates and incubated at 37 °C for 10-14 days. On the day of the readout, plates were stained with MTT (mixed 1:1 with PBS, 200 µl per 35 mm plate). MTT treated plates were incubated for 3-4 hours at 37 °C and visualized using the GE ImageQuant™ LAS 4000.

### Xenograft transplantation

The *in vivo* study carried out was approved by the Dana-Farber Cancer Institute Animal Care and Use Committee (IACUC). 7-week-old NOD/SCID/IL2rγnull mice (The Jackson Laboratory) were intravenously injected with 250,000 MV4-11 cells carrying Cas9 and sgRNA GFP inducible guides. After validation of engraftment via bone marrow aspiration, mice were started on a doxycycline (625 ppm) diet. Beginning day 4 of the doxycycline diet, mice were treated with GSK-3685032 (45 mg/kg/dose) or with vehicle (10 % captisol) via subcutaneous injection bi-daily for 7 days. Mice were sacrificed following the 7-day treatment to assess tumour burden. The leukemic burden was determined by flow cytometry analysis of mouse CD45-APC and human CD45-V450 staining of the peripheral blood, spleen, and bone marrow. The remaining mice were monitored for survival and time for sacrifice was determined by assessing body weight, overall fitness and signs of paralysis per DFCI IACUC protocol.

### Statistics and data analysis

#### Ubiquitinome data analysis

The normalized TMT data for sgNT and sgUSP48 guide #1 and #2 was provided for two time points, day 3 and day 7, in MV4-11 cells with two replicates per condition in two independent experiments. The TMT data was averaged across replicates and log2 transformed. Log2 fold change (LFC) intensity scores were computed for sgUSP48 versus sgNT. A cutoff of 0.5 for the absolute LFC and an adjusted p-value ≤ 0.10 for the limma eBayes test were used to assess significant changes. Data were visualized as hockey plots for differential *USP48* KO versus sgNT peptide changes induced by each guide and as gene-level LFC score heatmaps for selected paraspeckle and histone protein complexes in the Corum v3 database^54^. The gene-level LFC score per condition was computed as the maximum of the LFC intensity scores for that condition across the peptides mapped to the gene. A selected list of 340 proteins that scored with increased LFC TMT scores (≥ 0.50) for sgUSP48 versus sgNT at day 3 were further analysed for functional enrichment across Gene Ontology and Canonical Pathways (KEGG, Reactome) gene sets on the DAVID platform^55^. The top five identified functional clusters were visualized as an Enrichment Map network with gene sets represented as nodes and edges connecting two nodes if the gene sets overlap. Cutoffs for functional enrichment were set at FDR ≤ 0.05 for the hypergeometric test. Significance for gene set overlap was determined with FDR ≤ 0.05 for the hypergeometric test.

The TMT normalized data and the LFC intensity scores per condition are provided for each ubiquitinome assay in Extended Data Table 2, along with the functional cluster gene sets presented in the Enrichment Map network.

#### Proteomics

Proteome assay data was provided as mean TMT normalized intensity reporter scores for 4901 peptides associated to 4987 genes, for the same conditions and time points as the ubiquitinome data. The TMT scores were averaged per condition and log2 transformed. The differential changes induced by *USP48* KO vs. sgNT were evaluated based on the log2 fold change intensity scores with the cut-offs abs (LFC) ≥ 0.5 and adjusted p-value ≤ 0.10 for the limma eBayes test. Data were visualized as scatter dot plots of LFC intensity scores for *USP48* KO vs DMSO conditions. Similar to the ubiquitinome assay, the gene-level LFC score per condition was computed as the maximum of the LFC intensity scores for that condition across the peptides mapped to the gene. Extended Data Table 3 provides the TMT normalized proteome data and the LFC scores for the *USP48* KO vs sgNT conditions.

#### CoIP data analysis

The MV4-11 (L-IgG, H-USP48) CoIP assay yielded log2 H/L signal values for 410 peptides corresponding to 409 unique genes. LFC scores are provided in Extended Data Table 4. Interactor hits were identified using a threshold of log2 H/L ≥ 0.5. and relaxed interactor hits were identified using a threshold of log2 H/L ≥ 0.3. Gene set enrichment analysis for interactor hits was performed on CORUM v4.1 protein complex database^56^, and the Gene Ontology (c5) and Canonical Pathways (c2) collections from the MSigDB v4 database^57,58^. Statistical significance of overlap between the Co-IP hits and reference gene sets was evaluated using the hypergeometric test, with a significance threshold of adjusted p-value ≤ 0.05.

#### RNA-Seq data analysis

RNA-seq data were analysed in accordance with the ENCODE Consortium standards. Quality control of raw reads was performed using FastQC v0.12.1 (Babraham Bioinformatics) and summarized with MultiQC v1.9^59^. Reads were aligned to the hg38/Gencode v30 reference genome using STAR v2.7.2b^60^ with the standard parameters: --outSAMtype BAM SortedByCoordinate --outSAMunmapped None -- outSAMattributes NH HI NM MD AS XS --outReadsUnmapped Fastx -- outSAMstrandField intronMotif --quantMode TranscriptomeSAM GeneCounts --quantTranscriptomeBan IndelSoftclipSingleend. Post-alignment quality control and replicate reproducibility assessments were conducted using SARTools v1.7.3. Gene-level quantification was performed using featureCounts v1.6.3 from the Subread v2.0.0 package^61^, counting reads overlapping Gencode v30 gene exons. Gene expression was normalized and expressed as log2(TPM + 1). Genes with a maximum log2(TPM + 1) ≥ 1 across all conditions were considered expressed. Differential expression analysis was performed using the limma eBayes method^62^: genes were considered differentially expressed if they met both criteria: |fold change| ≥ 1.5 and adjusted p-value ≤ 0.10

#### Visualization and Functional Enrichment

Differentially expressed genes (DEGs) were analysed for functional enrichment using the DAVID platform^55^ across the following databases: Gene Ontology: Biological Process, Molecular Function, Cellular Component, and Canonical Pathways (Reactome, KEGG) databases. The top functional clusters enriched in the genes with increased expression by individual and combination treatment were assigned to four functional categories (1) Immune Response, (2) Chromatin Organization/Remodeling and Transcription Regulation, (3) DNA Damage and Repair (DDR) and (4) Cell death/Apoptosis. For each category, a consensus metagene was constructed by merging up to 10 representative gene sets from GO, Reactome, KEGG, and MSigDB Hallmark collections. The DDR metagene additionally incorporated the curated TCGA DDR gene set (n = 276 genes) from Knijnenburg et al.^63^. Gene set overlapping analyses for the DEGs vs. metagenes were performed with the significance cut-offs p-value ≤ 0.10, adjusted p-value ≤ 0.10 and size overlap ≥ 5 for the hypergeometric test. The enrichment scores computed as the percentage of overlapping (leadingedge) genes vs. the size of each metagene were visualized as dot plots. Enrichment plots for the metagenes vs. combination treatment conditions were visualized based on the GSEA software^57^, with the significance cut-offs: |Normalized Enrichment Score (NES)| ≥ 1.3, p-value ≤ 0.10, FDR ≤ 0.25. Heatmaps of transcriptional changes were generated using the Morpheus platform based on log2(TPM + 1) values and based on the log2 fold change scores. Differentially expressed genes per condition, selected metagenes, and enrichment results (DEGs vs. metagenes) are provided in Extended Data Table 5. Additional Enrichment Results: Enrichment analyses of DEGs vs. full Gene Ontology and curated pathway collections of gene sets are available upon request.

#### ATAC-Seq data analysis

ATAC-Seq data analysis was performed in alignment with the ENCODE Consortium standards (www.encodeproject.org/atac-seq/). Briefly, quality control for the pairedend unmapped reads was performed with the FastQC v.0.12.1 software (www.bioinformatics.babraham.ac.uk/projects/fastqc/). The reads were trimmed and filtered for Nextera adapters using fastp^64^. The trimmed reads were mapped to the hg38 reference genome using bowtie2-2.5.4^65^ with the –local –very sensitive –X 2000 options. Bam files were deduplicated using sambamba v1.0.1^66^. Only reads mapping to chromosomes 1 to 22 and chrX with an MAPQ of ≥ 5 were retained. Reads were shifted with 4 bp on the positive strand and −5 bp on the negative strand using AlignmentSieve available in deepTools v3.5.1^67^. Replicate correlations were computed with the multiBamSummary and bamCorrelate tools available in deepTools v3.5.1 and visualized as dendrograms and in principal components analysis plots. Fragment size distributions were computed with the bamPEFrag mentSize tool available in deepTools v3.5.1 and then inspected for quality control. Properly aligned reads for replicates were merged for each condition. Peak calling was performed for properly aligned and for the merged reads with MACS2 v2.1.1.20160309 software^68^ with the –nomodel, - - extsize to fragment length and –q 0.01 options. AUC binding signal for peaks was computed with the bwtool software^69^. The peaks were annotated with the closest hg38 gene TSS using the annotatePeaks function implemented in the Homer v4.11 platform^70^. Gene promoter regions were defined as the ±3.0 kb intervals around the hg38 gene TSS. Genome track bigwig files for normalized ATAC-Seq signal were created with the bamCoverage tool available in deepTools v3.5.1 with the Reads Per Kilobase per Million mapped reads (RPKM) method. Genome-wide ATAC signal was visualized as tornado plot heatmaps created with the computeMatrix, plotHeatmap and plotProfile tools available in deepTools v3.5.1. and on signal tracks with the IGV v2.19.2 platform^71^. The set of peaks were curated by removing regions overlapping with the ENCODE black-listed regions for hg38 (available at www.encodeproject.org/annotations/ENCSR636HFF/) and by filtering out peaks with low AUC signal. For each condition, the peak sites identified by MACS2 on merged reads were required the log2(AUC+1) signal of ≥10 for each replicate and log2 (AUC+1) ≥ 13.5 for the merged peak signal. Genome-wide nucleosome free positions were identified with the NucleoATAC tool^72^. A peak by sample counts matrix was created by counting the reads overlapping each aggregated peak with the multiBamSummary tool available from deepTools v3.5.1. Differential peak analysis for treatment vs sgNT conditions was performed on the merged peaks. For a specific genomic region, the changes in signal between two conditions were classified as increase, decrease or not significant on the basis of the absolute cutoff of 0.5 for the delta log2 (AUC+1) signal score and the cutoff P ≤ 0.10 for the differences in the mapped reads by using DiffBind edgeR v3.36.0 with the glmLRT model^73^.

Associations between changes in ATAC-Seq peak signals versus expression changes in the nearest annotated genes, induced by treatment compared to control, were visualized using scatter plots. Each dot in the scatter plot represents a gene. The merged ATAC peaks present in both treatment and control conditions were annotated with the nearest hg38 gene transcription start site (TSS) using Homer v4.11. This annotation process allows multiple ATAC-Seq peak regions to be associated with the same gene. For each annotated gene, the ATAC-Seq region closest to the gene’s TSS was selected. The scatter plot displays the Delta log2(AUC+1) score for the nearest ATAC-Seq region on the y-axis versus the log2(fold change) expression for the corresponding gene on the x-axis. Correlation analysis was conducted to estimate the Pearson correlation coefficient between changes in ATAC-Seq signal and changes in gene expression. Additionally, overlapping analysis was performed using the two-tailed Fisher exact test to identify genes with significant increases or decreases in ATAC-Seq signal and expression.

#### WGBS data analysis

Prior to alignment, sequencing reads were trimmed using fastp (version 0.22.0)^64^ and the following parameters: -- adapter_sequence=AGATCGGAAGAGCACACGTCTGAACTCCAGTCA - - adapter_sequence_r2=AGATCGGAAGAGCGTCGTGTAGGGAAAGAGTGT --trim_front2=15. Trimming is required to remove bases that are added during the Adaptase reaction that could affect alignment and DNA methylation calling. Processed reads were aligned to the GRCh38 reference genome using methylCtools (version 1.0.0, https://github.com/hovestadt/methylCtools)^74^ and bwa mem (version 0.7.17, arXiv: 1303.3997v1) using default parameters. Over 98% of reads were aligned as proper pairs across samples. After marking of PCR duplicates using sambamba (version 0.8.1)^66^, genome-wide CpG methylation values were called using methylCtools using the --trimPE parameter. Methylation calls per CpG were aggregated across strands. Average genomic CpG coverage was ∼10-fold for each of the 8 samples (2 replicates per treatment).

To analyse global DNA methylation patterns, CpG beta-values (ranging from 0 to 1) were averaged in 100kb tiling windows and visualized in IGV using a color scale from blue (0, unmethylated) to white (0.5, hemi-methylated) to red (1, fully methylated). For all other analyses, regions for individual CpGs were extended halfway to the preceding and following CpG, resulting in a continuous DNA methylation track.

Processed tacks were visualized using tornado plot heatmaps across 10 bp bins, generated via the computeMatrix and plotHeatmap modules in deepTools v3.5.1, with the bwr Matplotlib color scheme applied. Methylation signal across ATAC peaks or CpG island regions was computed by averaging beta-values across CpG sites overlapping the regions. A threshold of 0.5 was used to classify CpG islands as methylated (beta-value ≥ 0.5, red) or unmethylated (beta-value < 0.5, blue).

## Data availablity

CRISPR screen data are available in Extended Data Table 1. The ATAC Seq data, RNA-Seq data and WGBS data generated in this study are available in the NCBI GEO database under their BioProject IDs respectively. The proteomics, ubiquitinomics and interactome data will be deposited to the Proteomics Identifications Database (PRIDE) and released upon publication of this manuscript. Additional data sets will be available from the lead contact upon reasonable request.

## Code availability

No custom code was developed in this study.

---

**Extended Data Figure 1:**
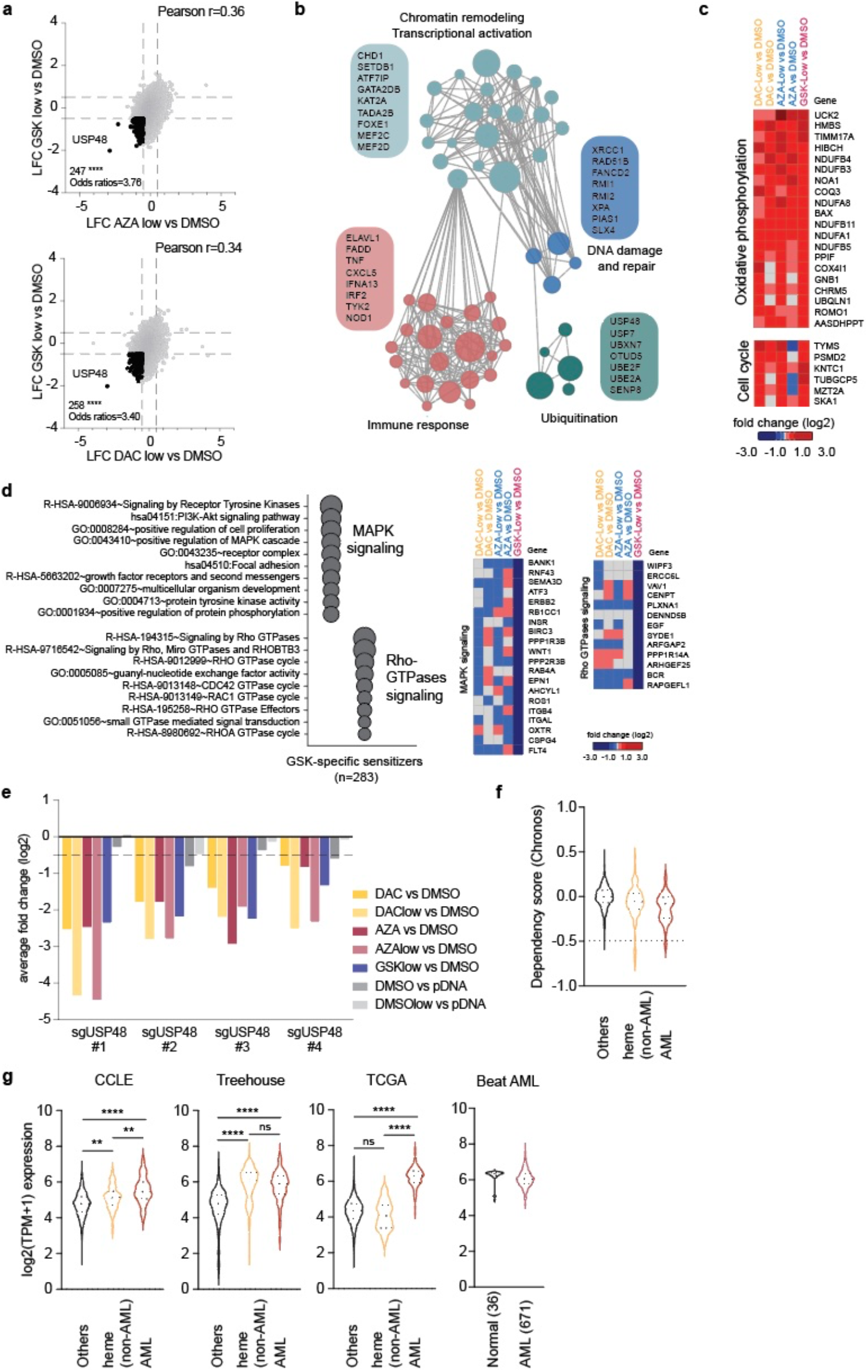
CRISPR screens identify USP48 as an HMA-sensitizer. **a)** Scatter dot plot showing the log2 fold change (LFC) of sgRNA abundance in **top)** AZA treatment vs. DMSO and GSK-3680532 (GSK) treatment vs. DMSO and **bottom)** DAC treatment vs. DMSO and GSK-3680532 (GSK) treatment vs. DMSO. The scores of all sgRNAs targeting the same gene were averaged and shown as one data point. Common consensus sensitizer genes are highlighted black. Significance cutoff: hypergeometric test abs (log2 fold change) ≥ 0.5, average p-value ≤ 0.1. Overlap significance: two-tailed Fisher exact test **** p < 0.0001. **b)** Enrichment Map network of the top functional clusters enriched in the 300 common consensus sensitizer genes for DAC, AZA and GSK in the CRISPR screen. Nodes represent gene sets, with node size indicating the number of genes in each set. Edges connect nodes (gene sets) that have a significant overlap (adjusted p-value ≤ 0.05 for the hypergeometric test). Shown in boxes are top leading-edge genes in each functional cluster. Functional enrichment analysis performed on the DAVID platform across Gene Ontology and Canonical Pathways (Reactome, KEGG) databases. Enrichment significance: adjusted p-value ≤ 0.05 for the hypergeometric test. **c)** Heatmaps depicting the CRISPR log2 fold change vs. DMSO scores for the top leading edge common resistance genes enriched in oxidative phosphorylation (left) and cell cycle (right). **d)** Bubble plot depicting the top functional clusters and gene sets enriched in the GSK-specific sensitizers **(left)**. Each bubble represents a gene set. The size of the bubble depicts the number of GSK-specific sensitizer genes in the associated gene set. Functional clustering analysis performed on the DAVID platform covering Gene Ontology and Canonical Pathways (Reactome, KEGG) databases. Enrichment significance: adjusted p-value ≤ 0.05 for the hypergeometric test. Heatmaps depicting the CRISPR log2 fold change vs. DMSO scores for the top leading-edge genes in the GSK-specific sensitizer functional clusters **(right). e)** Bar graph showing the log2 fold change (LFC) of sgRNA abundance of the four sgRNAs targeting *USP48* represented in the Avana library, for the DAC, AZA and GSK treated conditions and DMSO controls. Dashed black line at -0.5 indicates the gene effect score cutoff to be considered a dependency. **f)** Violin plots depicting the gene effect dependency scores for *USP48* in the 24Q4 DepMap CRISPR database across AML (n = 26, red), non-AML hematopoietic malignancies (n =104, orange) and solid tumour cell lines (n =1020, black). Dashed black line at -0.5 indicates the gene effect score cutoff to be considered a dependency. **g)** Violin plots depicting the *USP48* gene expression distribution across AML, non-AML hematopoietic malignancies and solid tumour samples in the 24Q4 CCLE, Treehouse, TCGA and Beat AML databases. One-way ANOVA with Tukey’s correction for multiple comparisons **** p < 0.0001, *** p < 0.001, ** p < 0.01, ns = not significant.

**Extended Data Figure 2:**
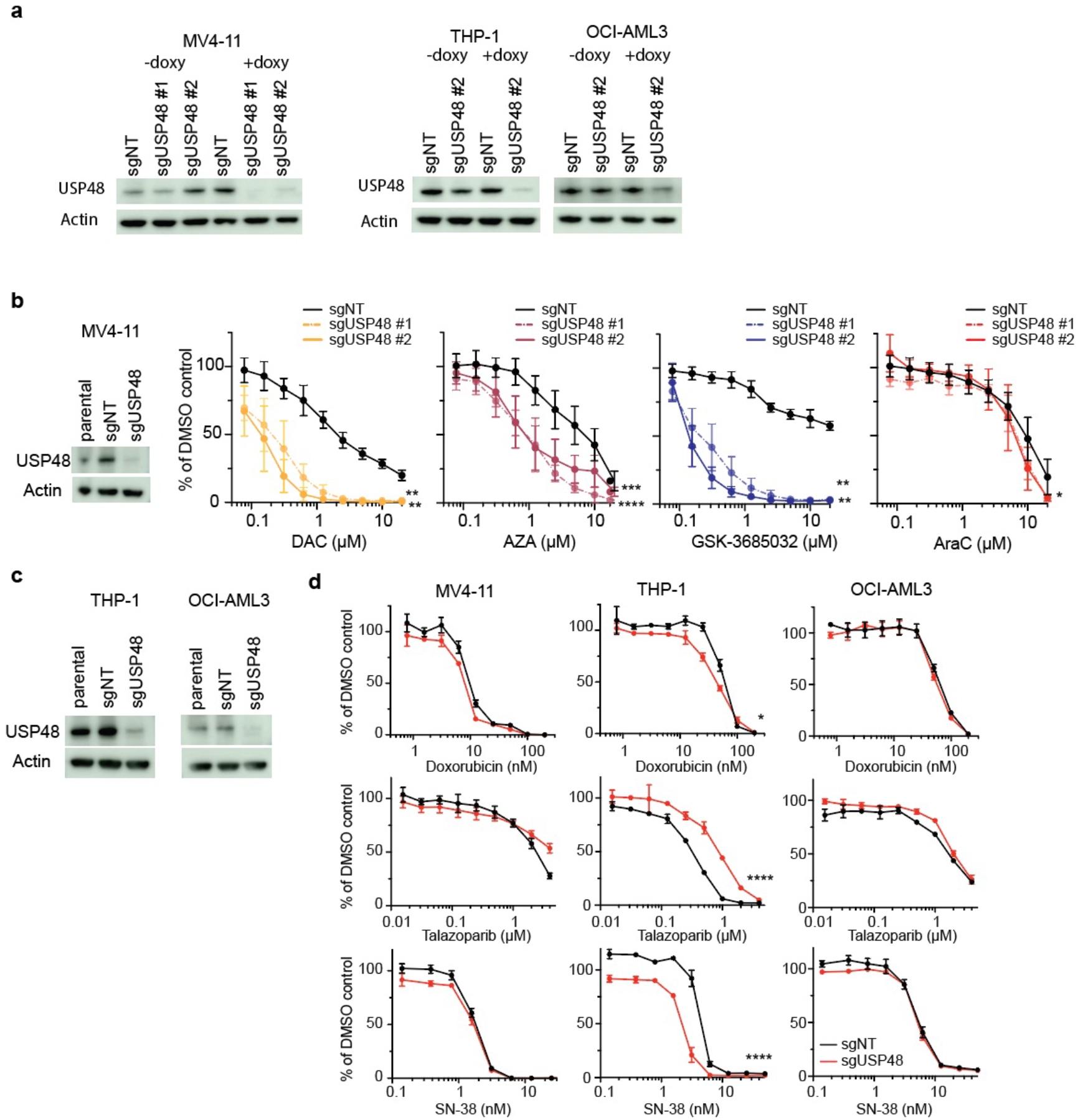
USP48 is an HMA-specific sensitizer. **a)** Validation of *USP48* KO after 4 days of doxycycline induction in MV4-11, THP-1 and OCI-AML3 in comparison to noninduced cells. Actin served as a loading control. **b)** Dose dependency curves for decitabine (DAC, yellow), azacytidine (AZA, burgundy), GSK-3685032 (blue) and cytarabine (AraC, red) measured after 96h of treatment in MV4-11 cells carrying control guides (sgNT, black) or guides targeting USP48, (sgUSP48 #1, dashed coloured line and sgUSP48 #2, coloured line). p-values were determined for extra sum of squares F-test LogIC50 differential best fit between pair-wise conditions **** p < 0.0001 *** p < 0.001, ** p <0.01, * p < 0.05. **c)** Validation of *USP48* KO in THP-1 and OCI-AML3 cells after 4 days of doxycycline induction using western blot. Actin served as a loading control. **d)** Dose response curves for MV4-11, THP-1 and OCI-AML3 sgNT (black) and sgUSP48 (red) cells after 96h of treatment with doxorubicin, talazoparib or SN-38. p-values were determined for extra sum of squares F-test LogIC50 differential best fit between pair-wise conditions * p < 0.05, **** p < 0.0001.

**Extended Data Figure 3:**
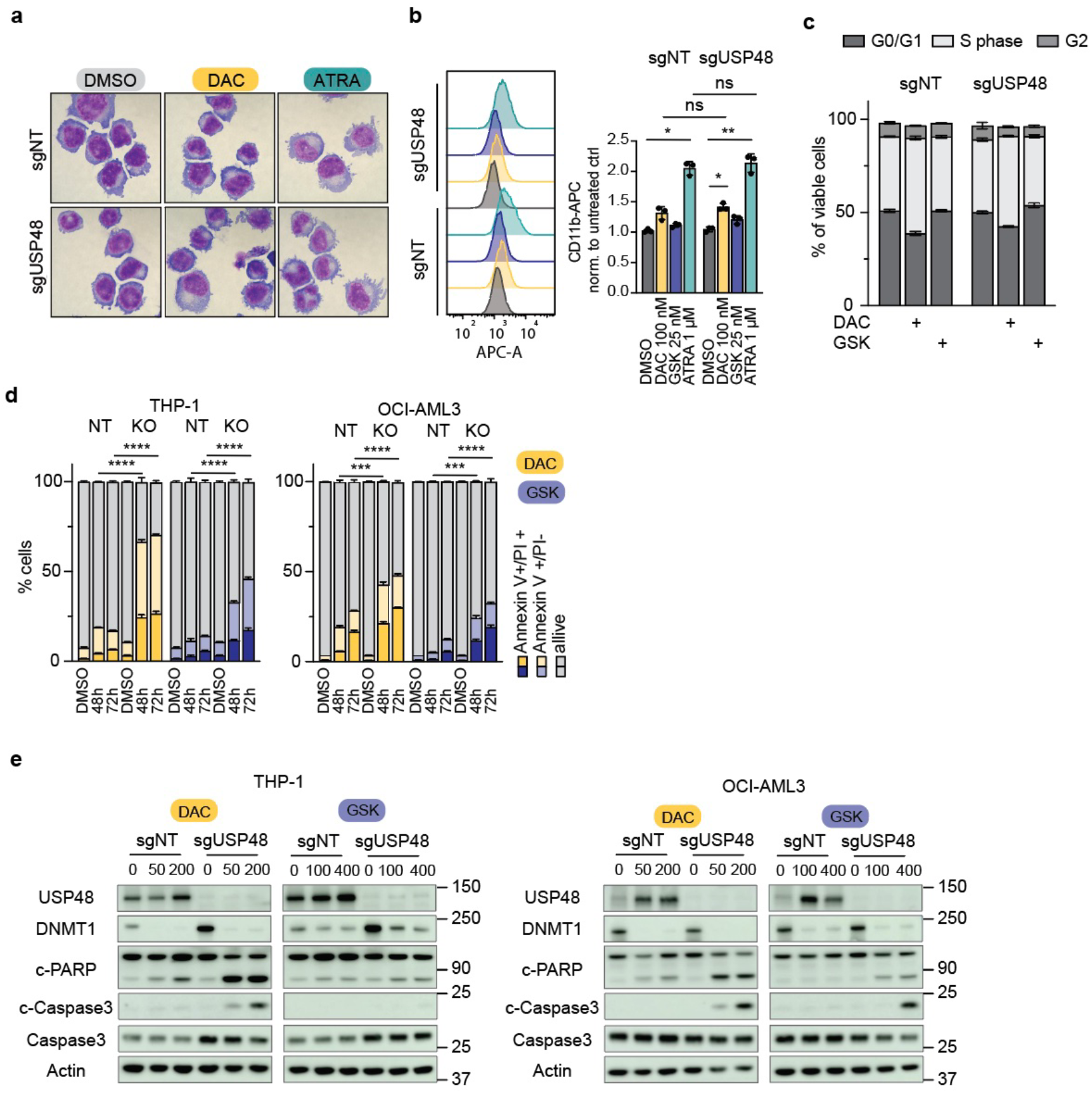
HMA treatment induces cell death in USP48 KO cells. **a)** Representative images of MV4-11 cells transduced with sgNT or sgUSP48 and treated for 72 h with 100 nM DAC or 1 µM ATRA. Cells were fixed, stained using May-Grünwald/Giemsa stain and morphology was assessed using a digital microscope. **b)** Flow cytometry analysis of CD11b expression in MV4-11 cells carrying sgNT or sgUSP48 after 72h of treatment with 100 nM DAC (yellow), 25 nM GSK (blue) or 1 µM ATRA (teal). Shown is a representative histogram for CD11b-APC signal and normalized CD11b expression to the untreated control (n=3). 2-way ANOVA multiple comparisons test with Tukey’s corrections was used to compare different drug treatment conditions in sgUSP48 vs. sgNT * p < 0.05, ** p < 0.01, ns-not significant. **c)** Cell cycle analysis of MV4-11 cells transduced with sgNT or sgUSP48 after 48h of treatment with 100 nM DAC or 25 nM GSK-3685032. Shown are percentages of viable cells in G2, S-phase and G0/G1 arrest. **d)** Annexin V/PI staining in THP-1 and OCI-AML3 sgNT and sgUSP48 cells treated with 200 nM DAC (yellow) or 400 nM GSK-3680532 (blue) for 48 and 72 h. Percentage of alive, early apoptotic (Annexin V+/PI-) and late apoptotic cells (Annexin V+/PI+) are shown per condition (n=3). One-way ANOVA multiple comparisons test with Tukey’s correction was used to compare sgUSP48 treated vs. sgNT treated conditions per time point *** p < 0.001, **** p < 0.0001. **e)** Western blots of THP-1 and OCI-AML3 sgNT and sgUSP48 cells treated with multiple concentrations of DAC or GSK-3680532 for 48h. *USP48* KO is validated and expression of DNMT1, cleaved PARP and cleaved Caspase 3 is shown. Actin served as a loading control.

**Extended Data Figure 4:**
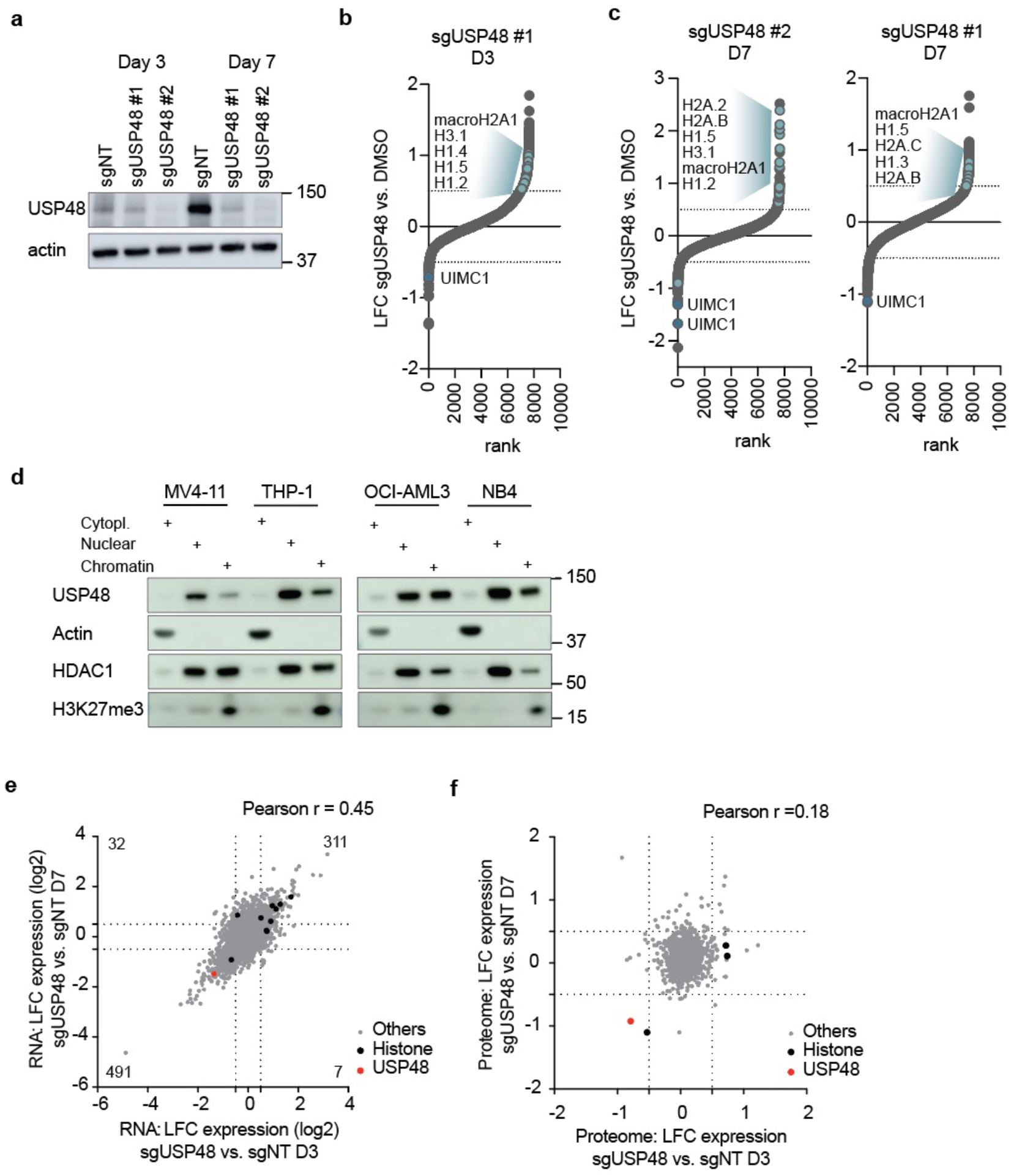
USP48 is localized in the nucleus and affects histone ubiquitination. **a)** Validation of *USP48* KO in MV4-11 after 3 and 7 days of doxycycline induction, using western blot, in the samples submitted for ubiquitinome and proteome analysis. Actin served as a loading control. **b and c)** Hockey plots of ubiquitinome data in MV4-11 after induction of *USP48* KO using **b)** sgUSP48 guide #1 after 3 days and **c)** sgUSP48 guide #1 or sgUSP48 #2 after 7 days. Each dot represents the log2 fold change (LFC) of one ubiquitin site identified using mass spectrometry analysis. Significantly increased or decreased histone ubiquitin sites (abs (LFC) ≥ 0.5) are highlighted in light blue. **d)** Western blot analysis of USP48 in cell fractionation samples from MV4-11, THP-1, OCI-AML3 and NB4 cells. Actin served as a marker for the cytoplasmic fraction. HDAC1 served as a marker for the nuclear and chromatin fractions and H3K27me3 served as a marker for the chromatin only fraction. **e)** Scatter dot plot of gene expression data in MV4-11 upon *USP48* KO after 3 and 7 days. Each dot represents the log2 fold change (LFC) of one gene. Significantly increased or decreased histone (abs (LFC) ≥ 0.5) are highlighted in black, USP48 is highlighted in red. Quadrant numbers represent overlap size. **f)** Scatter dot plot of proteome data in MV4-11 upon *USP48* KO after 3 and 7 days. Each dot represents the log2 fold change (LFC) of one protein identified using mass spectrometry analysis. Significantly increased or decreased histone (abs (LFC) ≥ 0.5 and adjusted p-value ≤ 0.10) are highlighted in black, USP48 is highlighted in re

**Extended Data Figure 5:**
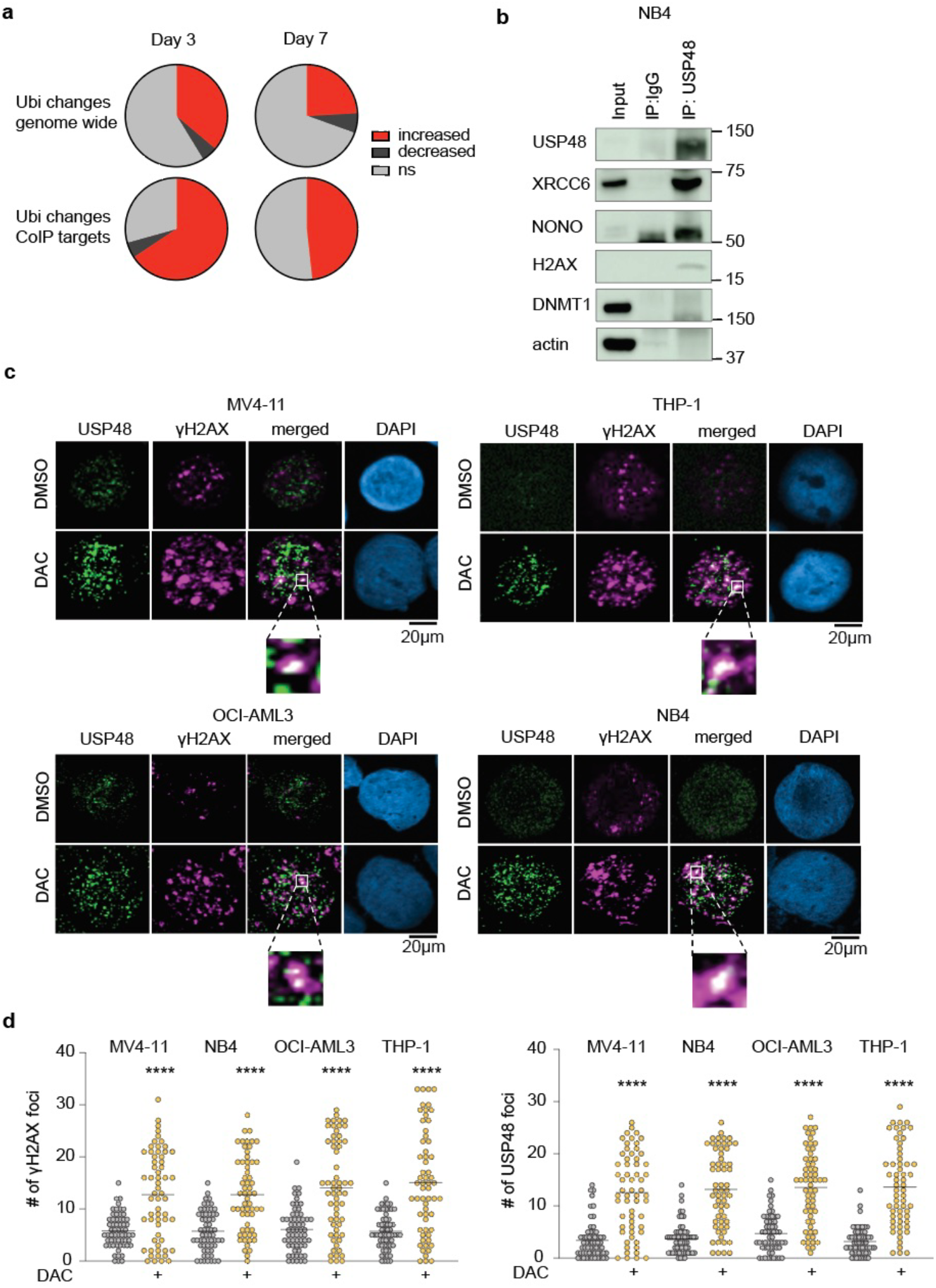
USP48 co-localizes with chromatin associated proteins and DNA damage sites. **a)** Pie-charts describing the ubiquitin changes (% proteins) induced by sgUSP48 vs sgNT at days 3 and 7 in the UBI2 ubiquitinome data set genome-wide and for the 67 Co-IP hits in MV4-11 cells. Significance: eBayes limma abs (fold change) ≥ 1.5, adjusted p-value ≤ 0.10. **b)** Western blot validation of USP48 co-immunoprecipitated proteins in NB4 cells in comparison to the input and the immunoprecipitated IgG control. **c)** Representative images of USP48-γH2AX foci in MV4-11, THP-1, OCI-AML3 and NB4 cells upon 48h DMSO or decitabine treatment (200 nM). Shown is staining for USP48 and γH2AX as single and merged images. DAPI is used for nuclear staining. **d)** Quantification of **left)** γH2AX foci and **right)** USP48 foci in MV4-11, NB4, OCI-AML3 and THP-1 cells after 48h treatment with 200 nM decitabine (DAC). Each dot represents one nucleus (n=60 nuclei per sample). 2-way ANOVA multiple comparisons test with Tukey’s corrections was used to compare DAC treatment conditions vs. DMSO per cell line, **** p < 0.0001.

**Extended Data Figure 6:**
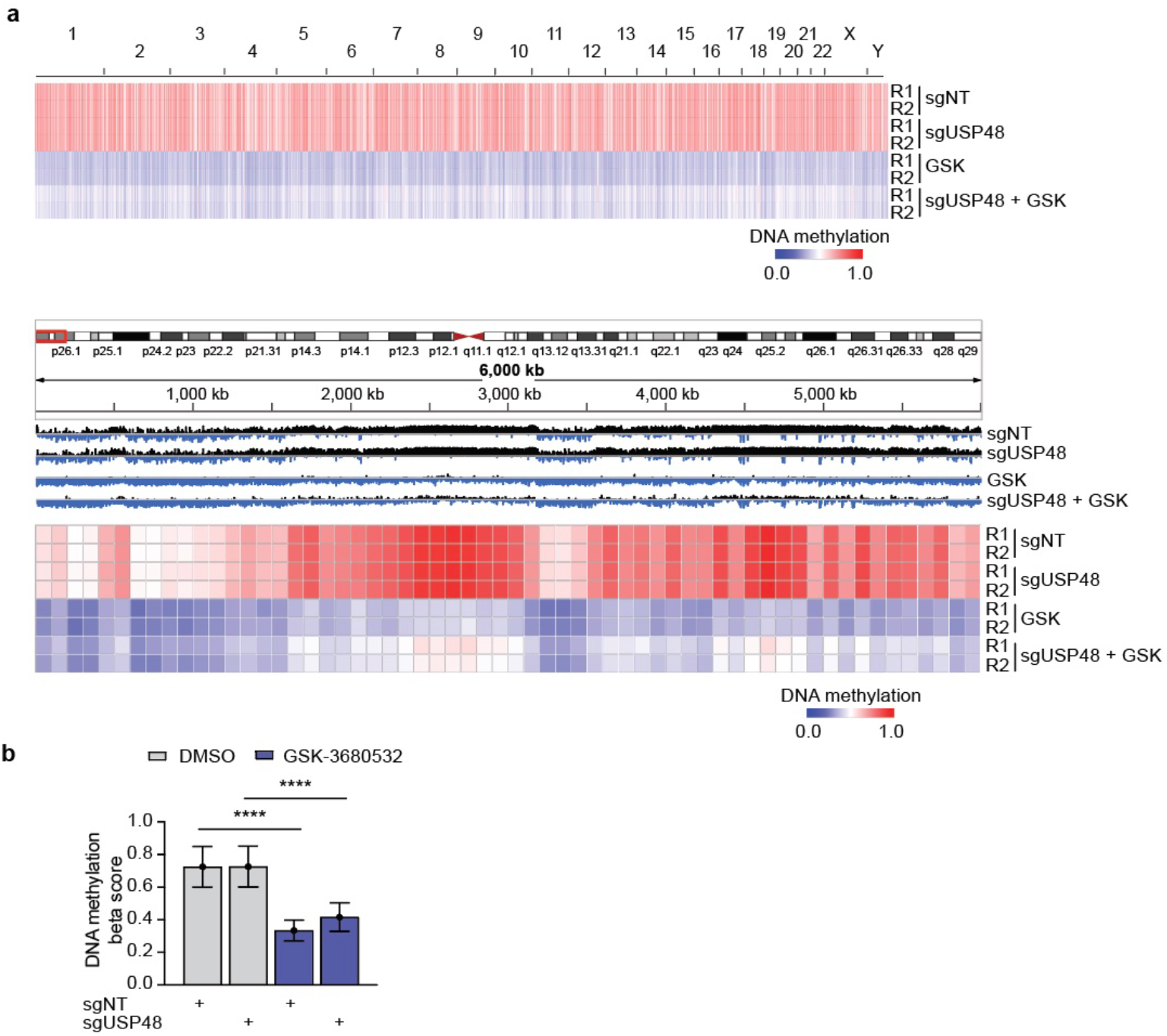
GSK-3680532 treatment results in genome-wide demethylation. **a)** Heatmaps presenting DNA methylation beta scores in 100 kb windows across various replicate conditions: sgNT, sgUSP48, GSK-3680532 (GSK), and sgUSP48 + GSK-3680532. Top: Genome-wide view. Bottom: Chromosome 3, range 1-6,000,000 bp. The IGV DNA methylation halfway beta score signal is displayed with the following data range: Min: 0.0, Mid: 0.5, Max: 1.0. Cutoffs: Hypomethylation: ≤ 0.5 (shown as IGV blue, heatmap blue), Hypermethylation: ≥ 0.5 (shown as IGV black, heatmap red). **b)** Mean with SD barplots for the genome-wide DNA methylation beta scores in 100 kb windows across conditions. One-way ANOVA with Tukey’s correction for multiple comparisons **** p < 0.0001.

**Extended Data Figure 7:**
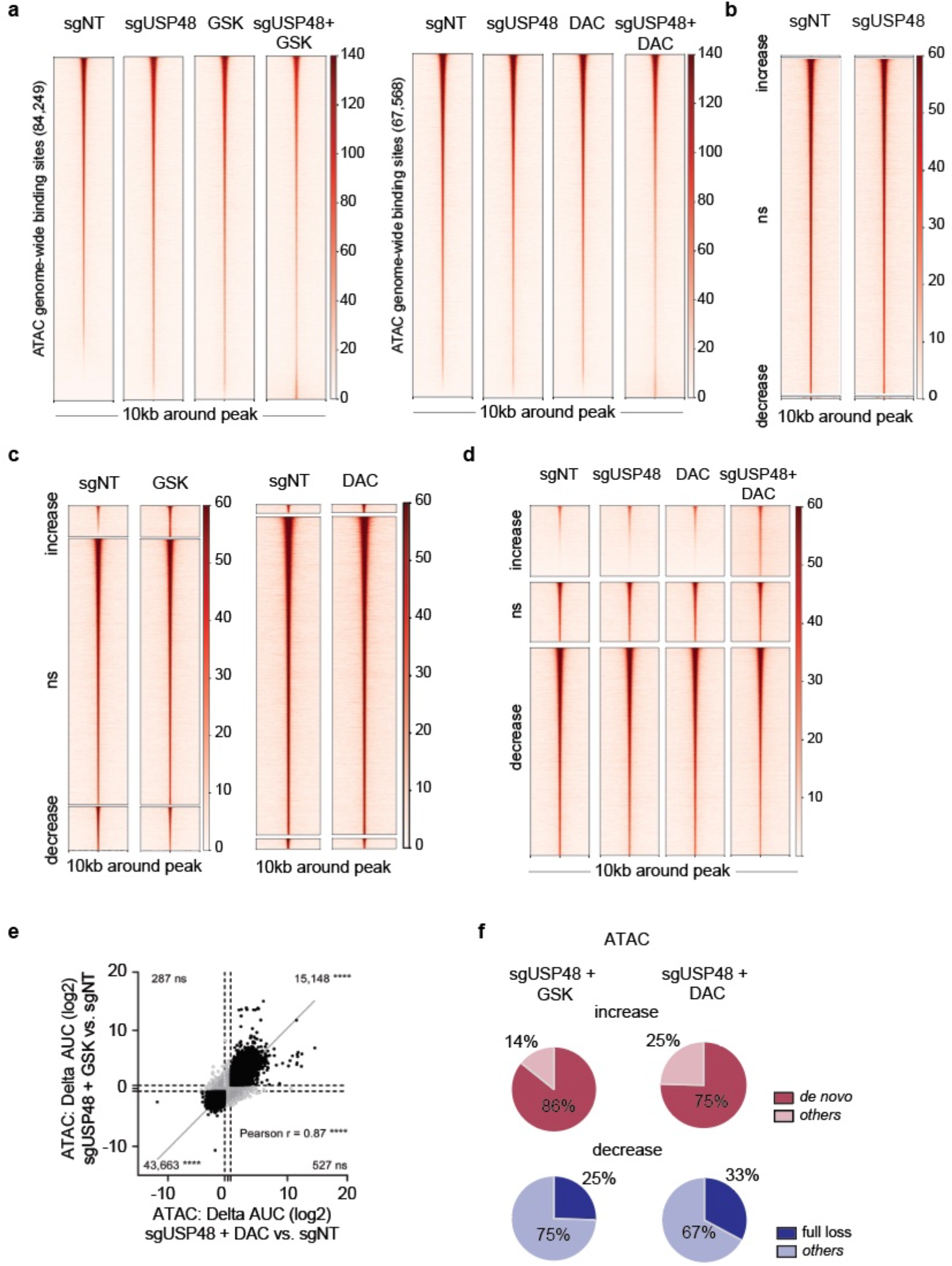
*USP48* KO increases chromatin accessibility upon DAC and GSK-3680532 treatment. a) Tornado plots depicting the ATAC signal on genome-wide chromatin accessibility regions across the treatment conditions **Left)** sgNT, sgUSP48, GSK, and sgUSP48 + GSK, **Right)** sgNT, sgUSP48, DAC, and sgUSP48 + DAC. The ATAC signal is shown as RPKM normalized. ATAC peaks are ranked by the ATAC AUC signal in the sgNT condition. **b)** Clustered tornado plots for RPKM normalized ATAC signal for sgNT vs. sgUSP48 in regions with increased, not significantly changed, and decreased AUC by sgUSP48 vs. sgNT. The cutoffs for differential signal are set at abs (Δ log2 AUC signal) ≥ 0.5, with a p-value ≤ 0.10. **c)** Clustered tornado plots for RPKM normalized ATAC signal in regions with increased, not significantly changed, and decreased AUC by **left)** sgNT vs. GSK-3685032 (GSK) and by **right)** sgNT vs. DAC. The cutoff for differential binding is set at abs (Δ log2 AUC signal) ≥ 0.5, with a p-value ≤ 0.10. **d**) Clustered tornado plots for RPKM normalized ATAC signal in regions with increased, not significantly changed, and decreased AUC by sgNT vs. sgUSP48 + DAC. The cutoff for differential binding is set at abs (Δ log2 AUC signal) ≥ 0.5, with a p-value ≤ 0.10. **e)** Scatter dot plot showing the correlation between the ATAC differential signal induced by sgUSP48 + DAC vs. sgNT and by sgUSP48 + GSK vs. sgNT. The cutoff for differential binding is abs (Δ log2 AUC signal) ≥ 0.5, p-value ≤ 0.10. The number of overlapping regions with significant differential signal for DAC and GSK are shown in the quadrants. Overlap significance was estimated using a two-tailed Fisher exact test (**** p < 0.0001, ns = not significant). **f)** Pie charts depicting the status of ATAC peaks with differential ATAC signal **left**) sgUSP48 + GSK vs. sgNT, and **right**) sgUSP48 + DAC vs. sgNT. These charts focus on regions with significant changes in ATAC signal induced by combination treatment vs. sgNT shown in the cluster in Fig. 3a and b **Top**: Fraction of *de novo* and other increased ATAC peaks in sgUSP48 + GSK or DAC treatment vs. sgNT. **Bottom**: Fraction of ATAC peaks in the sgNT condition that are lost or decreased in combination treatment.

**Extended Data Figure 8:**
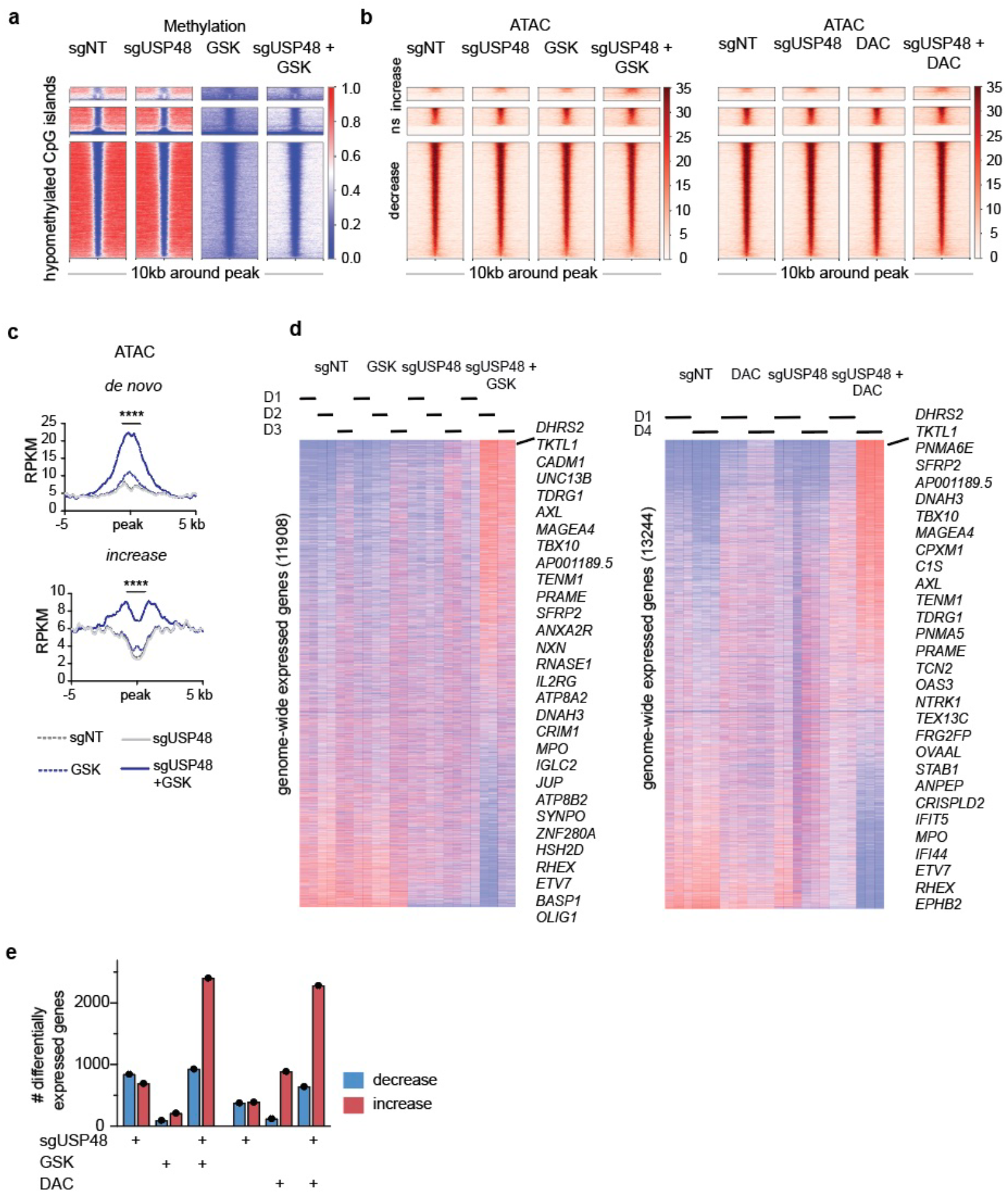
Combination of USP48 KO and HMA treatment leads to transcriptional activation. **a)** Clustered tornado plots illustrating the DNA methylation signals for sgNT, sgUSP48, GSK, and sgUSP48 + GSK treated MV4-11 cells. Shown are the genome-wide CpG island regions that are hypomethylated in the sgNT condition. Clusters display hypomethylated CpG islands with differential changes in ATAC signal induced by sgUSP48 + GSK vs. sgNT. Top: increased middle: not significantly changed, bottom: decreased. The cutoffs for differential signals are set at absolute change in log2 AUC signal (abs (Δ log2 AUC signal)) ≥ 0.5, p-value ≤ 0.10. DNA methylation is represented as halfway normalized beta scores. Scores below 0.5 indicate hypomethylation (blue), scores above 0.5 indicate hypermethylation (red). The hypomethylated CpG islands are ranked by ATAC sgUSP48+GSK signal within each cluster. **b)** Clustered Tornado plots depicting RPKM normalized ATAC signal across treatment conditions over the CpG islands shown at a). **Left**) sgNT, sgUSP48, GSK, sgUSP48+GSK and **right**) sgNT, sgUSP48, DAC, sgUSP48 + DAC. **c)** Metaplots illustrating the summary profiles of ATAC peaks on clustered hypermethylated CpG island regions from **Fig. 3e** for sgNT (light grey, dashed), sgUSP48 (grey), GSK (light blue, dashed), and sgUSP48 + GSK (blue). The plots display RPKM normalized scores for ATAC Seq. Differential signal was estimated using 1-way ANOVA with Tukey’s corrections for multiple comparisons (**** p < 0.0001). **d)** Heatmaps of genome-wide RNA-Seq expression data depicting genome-wide relative RNA-Seq expression across various replicate conditions and experiments. **Left:** sgNT, sgUSP48, GSK and sgUSP48 + GSK at days 1, 2, and 3. **Right:** sgNT, sgUSP48, DAC and sgUSP48 + DAC at days 1 and 4. The heatmaps are restricted to genes with maximum expression across conditions ≥ 1. Genes are ranked in decreasing order of sgUSP48 + treatment expression. **e)** Bar plots depicting the number of genes with increased and decreased expression induced by sgUSP48, DAC, GSK or sgUSP48 + DAC/GSK treatment vs. sgNT. Significance: eBayes (limma) abs (fold change) ≥1.5, adjusted p-value ≤ 0.10.

**Extended Data Figure 9:**
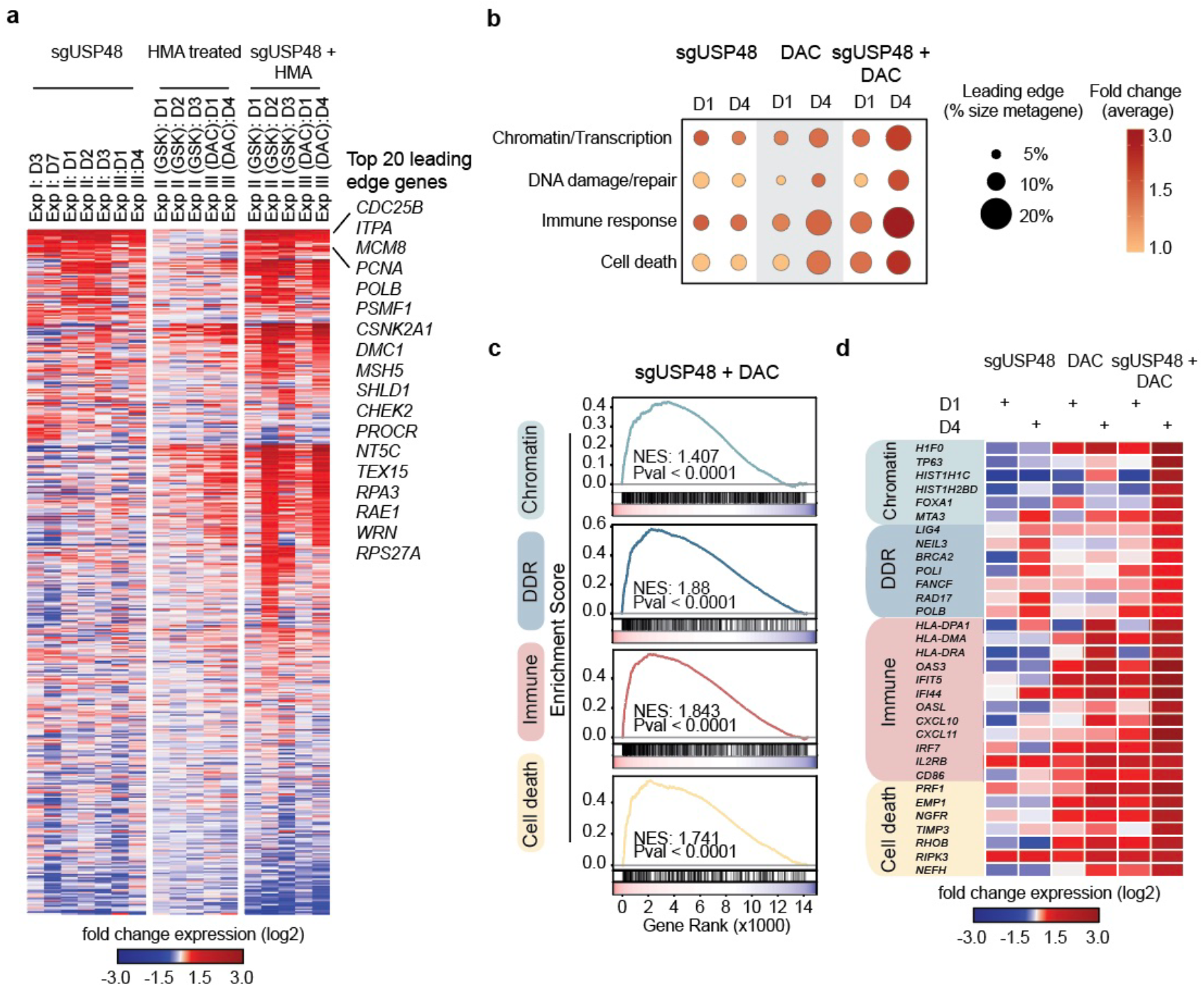
DNA damage and cell death pathways are activated upon HMA treatment in USP48 KO cells. **a)** Heatmap of changes in expression of DDR genes depicting the log2 fold changes in expression induced by *USP48* KO, GSK-3680532 (GSK) or DAC, and sgUSP48 + treatments (GSK or DAC) vs. sgNT across the DNA damage and repair (DDR) metagene in 3 independent experiments. The genes are ranked by the average expression change across all conditions. The top 20 DDR genes, which show a consistent increase in both sgUSP48 + treatment and sgUSP48 samples, are highlighted on the side. **b)** Bubble plot illustrating gene set enrichments associated with genes exhibiting increased expression in sgUSP48, DAC treated or sgUSP48 + DAC. Functional enrichment analysis was conducted using the DAVID platform, encompassing the Gene Ontology and Canonical Pathways (Reactome, KEGG) databases. The plot highlights the top four functional clusters, each represented by a metagene, created by merging the gene sets from the databases that describe the cluster. Bubble size indicates the percentage of genes in the metagene with increased expression. Colour reflects the average log2 fold change expression across the leading-edge genes. **c)** GSEA plots for the 4 metagene sets presented in **b)** for sgUSP48 + DAC at day 4. Significance abs (NES) ≥ 1.3, *P*-value ≤ 0.10, FDR ≤ 0.25. **d)** Heatmap of log2 fold expression changes of top leading-edge genes per functional cluster for sgUSP48, decitabine treatment alone (DAC) or in sgUSP48 + DAC at days 1, 2 and 3 of treatment.

**Extended Data Figure 10:**
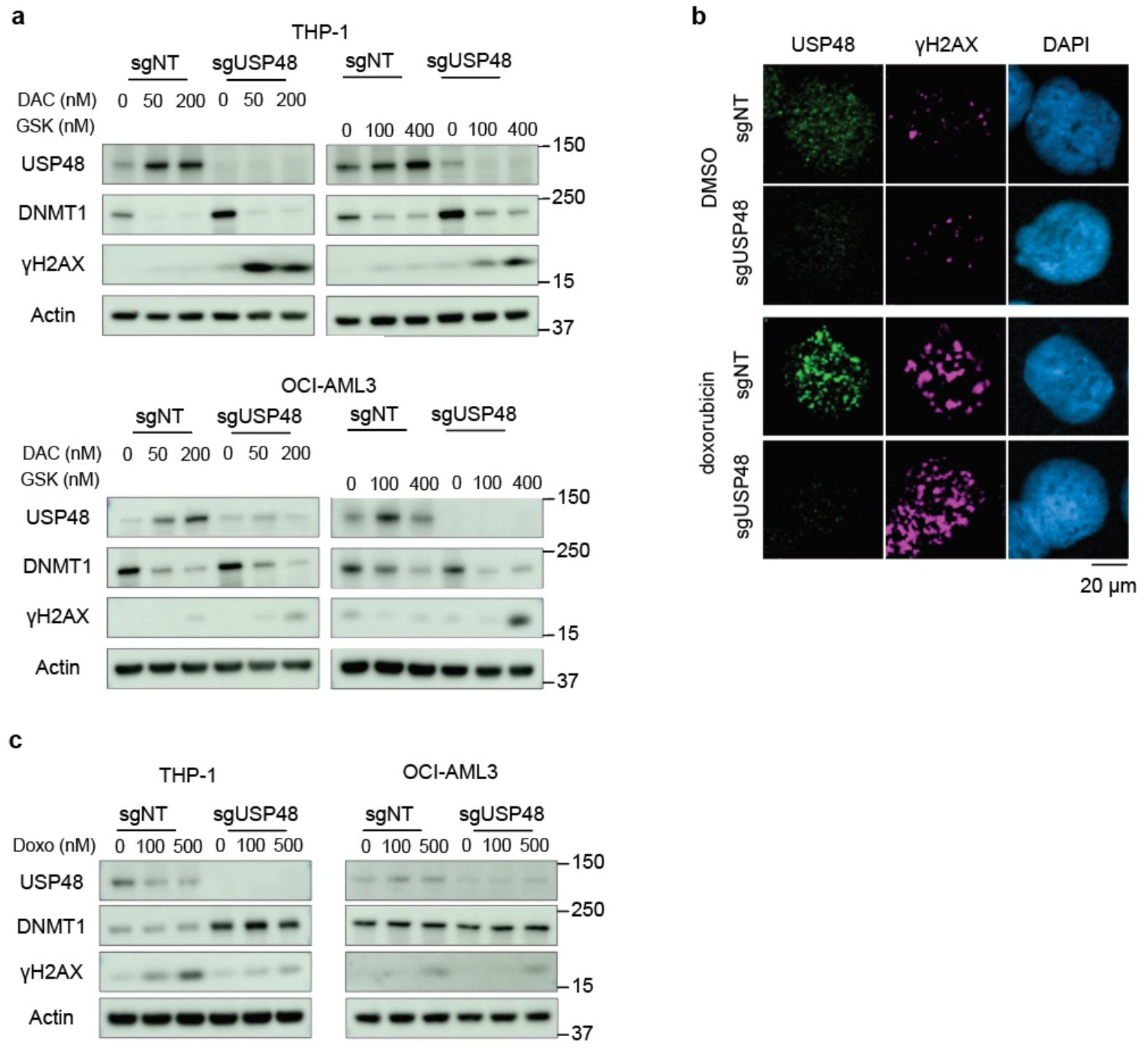
DNA damage phenotype in *USP48* KO cells is confirmed via western blot and immunofluorescence staining. **a)** Western blots of THP1 and OCI-AML3 sgNT and sgUSP48 cells upon 48h treatment with decitabine (DAC) or GSK-3680532 (GSK). *USP48* KO is validated and levels of DNMT1 and γH2AX are shown. Actin served as a loading control. **b)** Evaluation of USP48 and γH2AX antibodies used in immunofluorescence staining. Shown are representative images of MV4-11 sgNT and sgUSP48 cells upon 6h DMSO or doxorubicin treatment (500 nM). DAPI is used as nuclear staining. **c)** Western blots of THP1 and OCI-AML3 sgNT and sgUSP48 cells upon 6h treatment with doxorubicin. Antibodies for USP48, DNMT1 and γH2AX are shown. Actin served as a loading control.

**Extended Data Figure 11:**
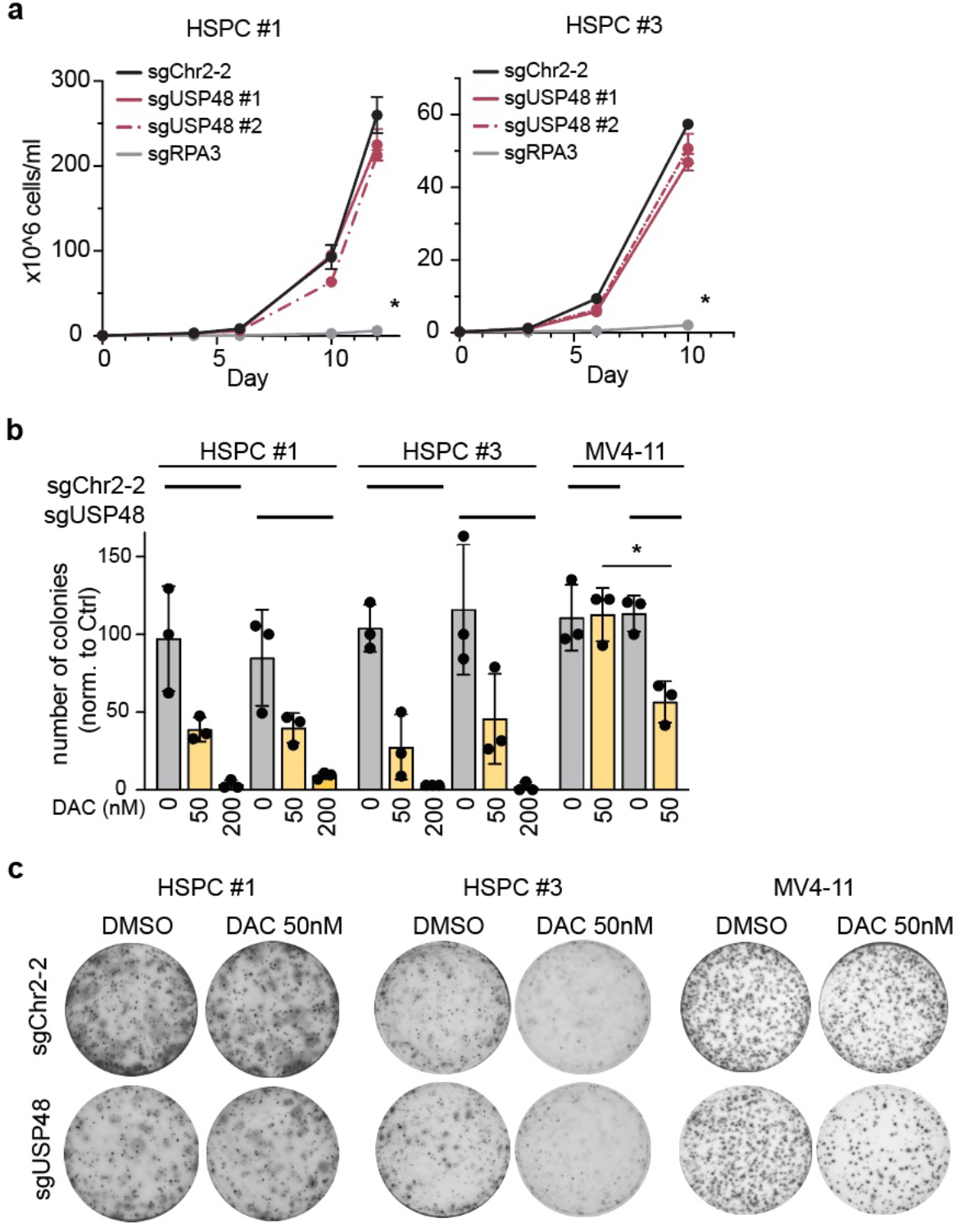
Viability of normal HSPCs is not affected by *USP48* loss. **a)** Counting experiment of HSPC sample #1 and #3 carrying sgUSP48 #1 (red), sgUSP48 #2 (red dashed), sgRPA3 (grey) or sgChr2-2 (black) over 10-12 days. One-way ANOVA multiple comparisons test with Tukey’s corrections was used to compare KO of *USP48* and *RPA3* vs. sgChr2-2 at Day 10-12. * p < 0.05. **b and c)** Colony formation assays of HSPC sample #1, HSPC sample #3 and MV4-11 cells nucleofected with sgChr2-2 or sgUSP48 #2 with or without decitabine (DAC) treatment for 10-14 days. At day of readout, plates were stained with MTT and visualized 3-4 hours later by microscopy imaging (n=3). Number of colonies was evaluated in triplicates from representative plates shown in **c)** for HSPC sample #1, HSPC sample #3 and MV4-11 cells nucleofected with sgChr2-2 or sgUSP48 #2. Cells were treated with DMSO (grey) or decitabine (DAC, yellow) for 10-14 days. Each dot represents one plate per condition (n=3); values are normalized to DMSO control.

**Extended Data Figure 12:**
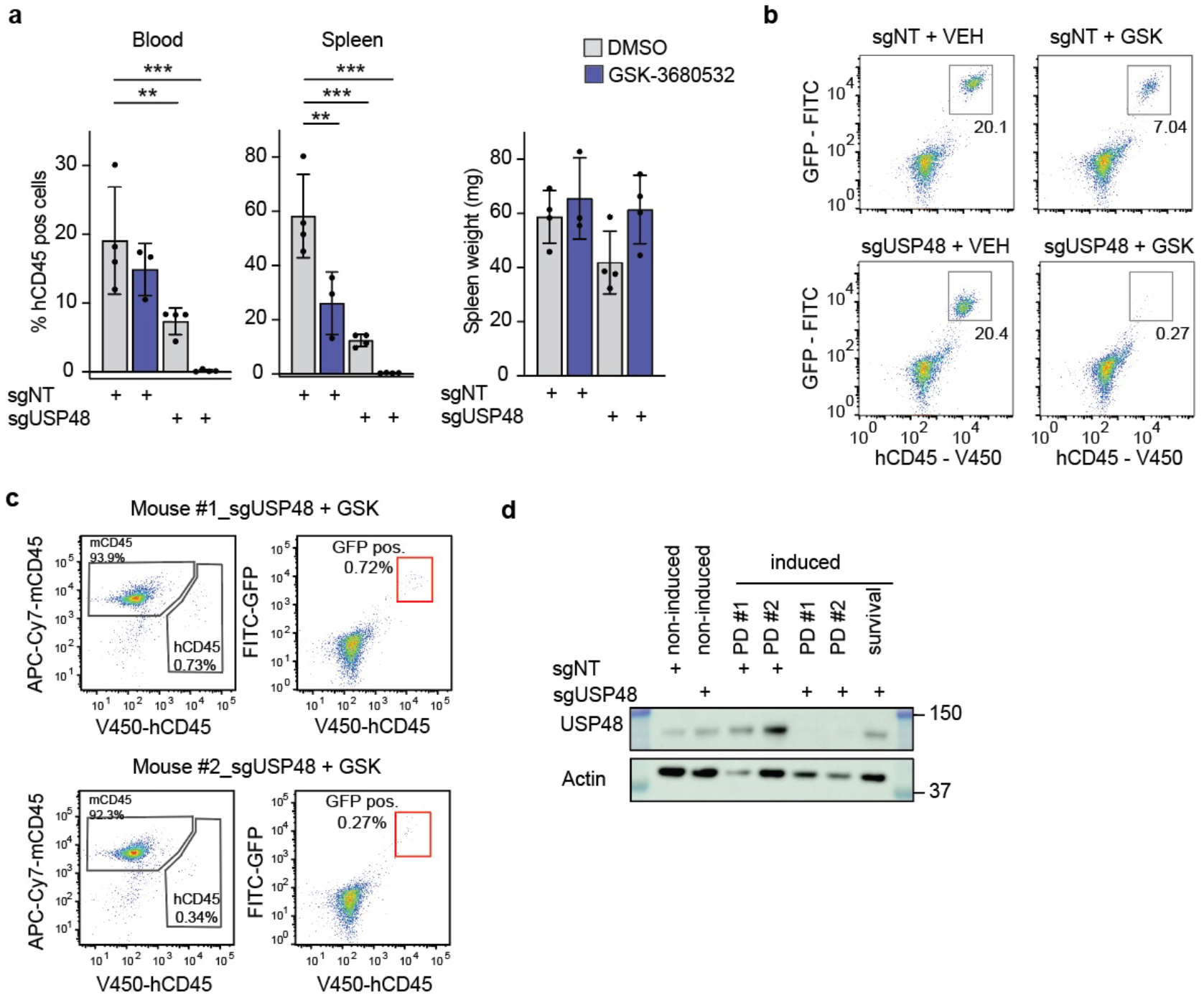
Combination of *USP48* KO and GSK-3680532 treatment ablates AML blasts *in vivo*. **a)** Left) Bar graph of % human CD45 positive cells in blood and spleen after 7 days of treatment with the vehicle control (DMSO, grey) or GSK-3685032 (GSK, blue). Each dot represents the measurement of one mouse. Right) Spleen weight of mice assessed after 7 days of treatment. Each dot represents the measurement of one mouse. Ordinary one-way ANOVA multiple comparisons test with Tukey’s correction was used to compare treatment conditions vs sgNT + VEH *\**\* p < 0.01, *** p < 0.001. **b)** Representative flow cytometry gating of bone marrow samples of sgNT and sgUSP48 mice treated with vehicle (VEH) or GSK-3680532 (GSK) assessed after 7 days of GSK treatment. Samples were stained for human CD45-V450, and GFP-FITC signal from the sgRNA vector was used to gate for MV4-11 cells. Percentage of GFP positive hCD45 cells is shown. **c)** Bone marrow samples of censored mice at time of sacrifice, stained for mouse CD45-APC-Cy7 and human CD45-V450. Percentage of mCD45 and hCD45 are shown. Red box highlights hCD45 positive MV4-11 sgNT-GFP and MV4-11 sgUSP48-GFP cells. **d)** Western blot of USP48 from samples of uninduced MV4-11 sgNT and sgUSP48 cells before injection, at time of PD study (10 days after doxycycline diet, 7 days after GSK-3680532 treatment was started) and in a sgUSP48 mouse monitored for survival at endpoint. Actin served as a loading control.

